# Control of timing and directionality during neurodifferentiation of human induced Pluripotent Stem Cells (iPSC) via miRNA-mediated feedback and feedforward loops

**DOI:** 10.1101/2023.10.16.562489

**Authors:** Jamie S Wood, Mouhamed Alsaqati, Jeremy Hall, Adrian J Harwood

## Abstract

Development of neural cells follows a unidirectional path, progressing from pluripotent stem cells into a variety of highly differentiated neuronal and glial cell types. Although the intermediate progenitor cells along this path are well characterised, the mechanisms that determine their directional flow remain to be established. Here, we identify a gene regulatory network that directs progression of neuronal differentiation. Previously, it was shown that loss of the epigenetic repressor, EHMT1, leads to premature neuronal differentiation, and this correlates with elevated miRNA expression and inhibition of a neuronal gene suppressor, REST. Here, we use a combination of *in silico*-analysis and targeted suppression of multiple miRNAs to reveal an unexpected degree of complexity to this gene regulatory pathway. We identity four miRNA, miR26a, miR140, miR142, miR153, that cooperatively block REST at the mRNA level. In contrast to most miRNA suppressors of REST, such as miR9a and miR124, which are in turn transcriptionally repressed by REST in a negative feedback loop (FBL), miR26a, miR140 and miR142 are not REST-targets. This creates a feedforward loop (FFL) that drives developing cells to switch from a high REST, non-neuronal state to a low REST, neuronal state, but not in the reverse direction. We show that together miR26a, miR140, miR142 and miR153 are necessary for neuronal differentiation and introduction of synthetic miRNAs mimics is sufficient to increase downstream miR-9 and miR-124 expression. Finally, we note that the Neurodevelopmental Disorder (NDD) Kleefstra Syndrome (KS) is caused by the loss of EHMT1, suggesting that the molecular mechanism described above may play a significant role in human brain development.

## Introduction

Neurodevelopment of the human brain occurs over an extended period, first during the prenatal phase of brain development and then via continued refinement to tune brain function over the following decades, stretching deep into adulthood. In contrast, we know that at the cellular level human induced pluripotent stem cells (iPSC) can be driven *in vitro* to form functional neurons in less than two weeks[1]. This argues against neuronal differentiation *per se* as a limit for brain development but rather a need to control when and where neural cell differentiation occurs. In fact, a highly organised temporal and spatial regulation of human neurodevelopment may be fundamental for the creation of the brain’s high-level functional properties[2]. Epigenetic gene regulation is instrumental in control of the timing of human neuronal development [3] but the mechanistic details by which it acts remain to be worked out. Here we demonstrate a complex miRNA-mediated gene regulatory feedback loop that de-represses neural gene expression to control when neurodevelopment can proceed.

The importance of epigenetic regulation for human brain development is underlined by its association with neurodevelopmental disorders (NDDs), where aberrant differentiation of neurons can lead to Autism (ASD), Schizophrenia (SCZ) and Intellectual disability (ID). In general, the genetic underpinnings of these conditions are highly polygenic and involve a combination of large numbers of risk loci, each of low effect [4–6]. However, a growing number of genetic syndromes have been discovered that individually impart high risk for NDDs, and many are associated with dysregulated epigenetic mechanisms [7–9], especially genes encoding histone methyltransferases and chromatin remodelers [10]. The number of known causative mutations has risen from 20 in 2014 to the most recent estimates identifying 179 disorders associated with 148 epigenes, and over 20% of all 720 epigenes causative for at least one chromatinopathy [11, 12]. Furthermore, it is estimated that nearly all chromatinopathies have some degree of neurological impact, with 85-93% displaying symptoms of intellectual disability [12, 13].

The gene EHMT1 encodes Euchromatic histone-lysine N-methyltransferase 1, a histone methyltransferase responsible for the H3K9me2 repressive chromatin mark. It is causative for Kleefstra Syndrome (KS), a NDD characterised by intellectual disability, psychotic disorder, microcephaly and facial dysmorphia [14]. EHMT1 is a key regulator of neurodevelopment, and microdeletions, loss of function mutations, and microduplications have been associated with NDDs [15, 16]. EHMT1 is required to regulate homeostatic plasticity via synaptic scaling; regulate neuronal excitability through excitatory-inhibitory balance; and control normal cranial development [17–19]. Recent work has demonstrated that reduced EHMT1 activity in KS-patient induced pluripotent stem cells (iPSC) results in up-regulation of microRNA (miRNA) expression and is associated with accelerated neuronal differentiation[20]. Amongst their many functions, miRNAs are essential regulators of every stage of brain development and maturation, and deregulation of some groups of miRNAs are associated with mental health conditions, such as Autism, Fragile X syndrome and Schizophrenia [21–23]. Although suggesting a direct link between EHMT1 activity and miRNA-mediated regulation during early stages of neurodevelopment, how this operates and impacts on neurodevelopment remains to be established

Here we identify a small subgroup of miRNAs that is elevated when EHMT1 activity is reduced and show that they control the master regulator REST and its brain-specific miRNA targets. This interaction creates a neurodevelopmental switch that in its initial state suppresses neuronal differentiation via presence of REST protein but then is rapidly and irreversibly lost to allow development to proceed. As a consequence, this switch controls the timing and sets the forward trajectory of neuronal differentiation towards neuronal maturity.

## Results

### EHMT1 activity and regulation of neuronal differentiation

We previously found that reduced EHMT1 activity and subsequent loss of its repressive histone mark H3K9me2 in pluripotent stem cells caused early and rapid neurodifferentiation [20]. To investigate the mechanism by which reduced epigenetic repression at the pluripotent stage leads to later altered neurodevelopmental timing, we examined changes in neurodevelopmental gene regulation during the neural progenitor stage. Fear and coworkers previously described REST-regulated genes in a EHMT1 hemizygous human mutant (EHMT1^+/−^) iPSC cell line following its differentiation to the Day 24 neural progenitor stage (NPC) [24]. We re-analysed their EHMT1^+/−^ NPC RNAseq data to identify 383 differentially expressed genes (DEGs) compared to wild type, with 231 upregulated and 151 downregulated (Fig. 1A). Thirty-six of the 231 upregulated DEGs (15.6%) were associated with a REST binding site (RE1); indicating an enrichment of REST-regulated genes amongst the set of upregulated DEGs (empirical p = 1×10⁻⁴ by permutation testing; Fig. S1).

**Figure 1:**
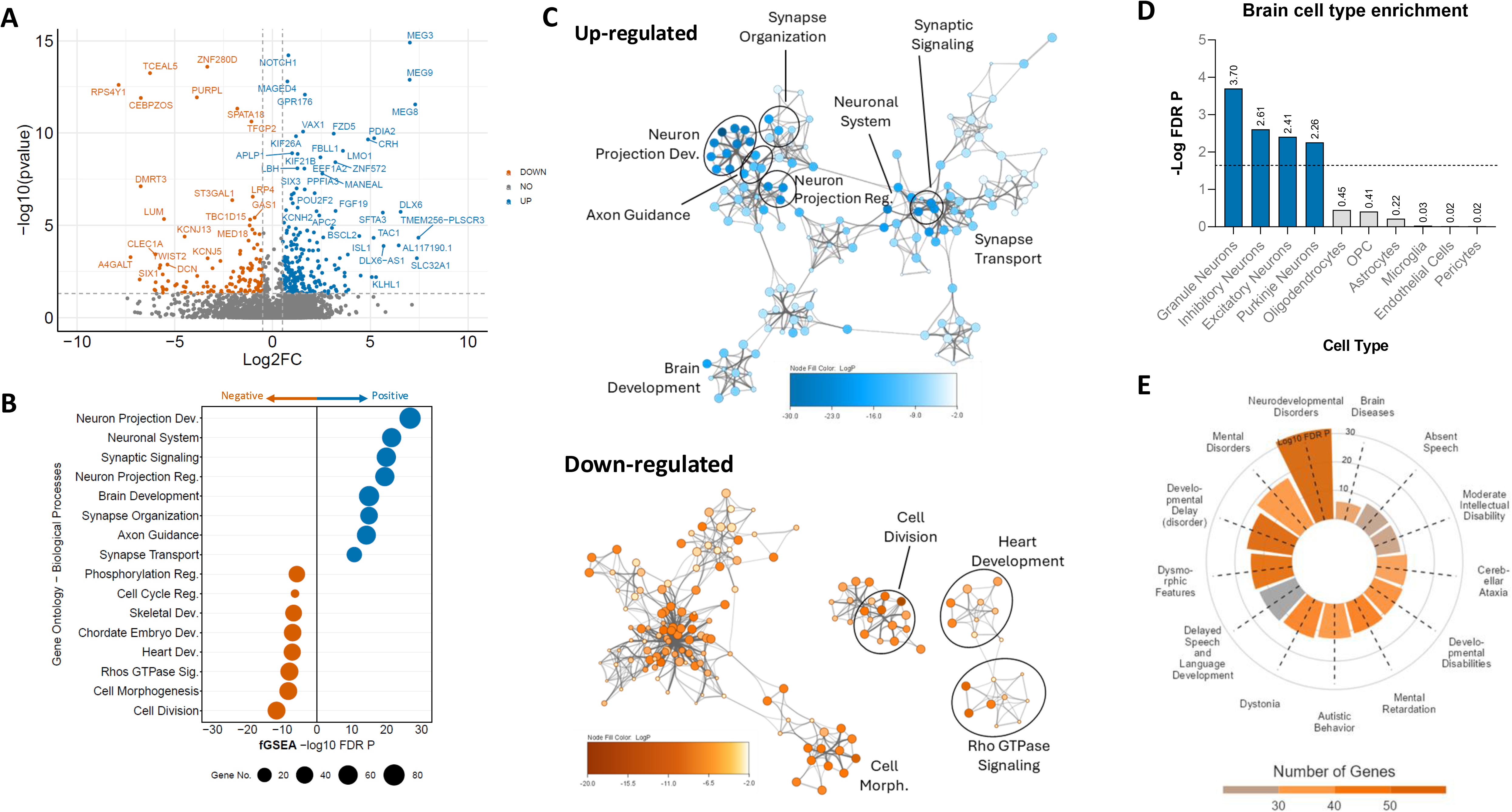
Loss of EHMT1 leads to increased neuronal differentiation: **A:** Volcano plot of differentially expressed genes (DEGs) in EHMT1^+/−^ human iPSCs versus wild type, analysed with DESeq2 (Wald test) using a log_2_ fold change (log_2_FC) threshold of ±1.5 (≈2.8-fold) and Benjamini–Hochberg FDR < 0.05. **B:** Gene set enrichment analysis (GOBP) on ranked genes (rank metric: signed minimum significant difference, msd), using fgsea multilevel; pathways significant at FDR < 0.05. **C:** Network of enriched GOBP terms, clustered by shared ID and coloured by p-value, for both upregulated DEGs and downregulated DEGs. Enriched GOBP terms were clustered based on semantic similarity using GOSemSim (R package, v2.24.0), and clusters were manually summarized. Node size indicates the number of genes in each term; node colour indicates adjusted P value (from fgsea). **D:** Brain cell type enrichment analysis using gene expression per tissue based on brain tissue single cell data. Significant cell types P<0.05, are in blue. **E** Disease-gene enrichment analysis performed for all DEGs and disease associations from DisGeNET, with significance determined as P < 0.001. Bar plot colour indicates the number of enriched genes within a given disease gene-set.

To select a relevant gene set for our further analysis, we identified significantly up and downregulated genes due reduced EHMT1 activity and performed Gene Set Enrichment Analysis (GSEA) based on the Gene Ontology (GO) database [25]. For upregulated genes, enriched GO Biological Processes (GOBP) are associated with neuronal development, including Neuron Projection Development, Synaptic Signalling, Brain Development and Synapse Organization (Fig. 1B & C). In contrast, downregulated genes are enriched for processes including Cell Cycle Regulation, Cell Morphogenesis and Cell Division (Fig. 1C). Further, tissue analysis indicated significant enrichment for brain specific gene expression, as well as ovary, heart and pancreas (Fig. S2), and brain cell type enrichment analysis based on single cell sequencing reference datasets from human neural tissue [26] indicated significant enrichment for all subtypes of neuron found within the brain including, granule, inhibitory, excitatory and cells (Fig. 1D). Finally, as the EHMT1 gene is causative for Kleefstra Syndrome (KS), we assessed gene-disease association using the *DisGeNET* database [27]. Of its 13 associated disease classes, Neurodevelopmental Disorders showed the highest enrichment, as well as the greatest DEG overlap (Fig. 1E). Interestingly, the remaining diseases and disorders showed significant overlap with other observed phenotypes for KS, including Dysmorphic Features, Delayed Speech and Language Development and Dystonia. Commonly enriched genes included NLGN3, KIF1A, EEF1A2 and MAST1, de-novo mutations for each gene being linked to a range of neurodevelopmental symptoms [28–32].

Collectively, these data further support our earlier observations that reduced EHMT1 activity alters the transcriptional programme of *in vitro* neurodevelopment by accelerating progression along the neuronal differentiation trajectory [20]. Furthermore, our refined DEG set shows significant enrichment for genes linked to NDDs with phenotypic overlap with Kleefstra syndrome, suggesting that related molecular programs may be perturbed *in vivo*.

### EHMT1 activity orchestrates the interplay between REST and miRNAs

We previously found that in pluripotent iPSCs (Day 0) reduced EHMT1 activity leads to upregulation of 56 miRNAs, of which 11 are predicted to target REST mRNA and cause its degradation or block its translation into protein [20]. We next applied an *in silico* prediction tool using a knowledge guided integrative analysis of miRNA-mRNA interactions, as previously described for analysis of ASD and pancreatic cancer associated miRNA [33, 34], and used it to identify potential EHMT1-sensitve miRNA regulators during early neurodevelopment (Day 24). This analysis identified 18 miRNAs that are likely to be upregulated in EHMT1 mutant NPCs at Day 24 (Fig. 2A). Eight of these predicted miRNAs have associated REST-binding (RE1) sites and been shown to be repressed by the REST complex [35]. These included the brain specific miRNAs miR-124 and miR-9, which are not only repressed by REST binding, also target REST or Co-REST mRNA to block REST action in a negative feedback loop [36] (Fig 2A).

**Figure 2:**
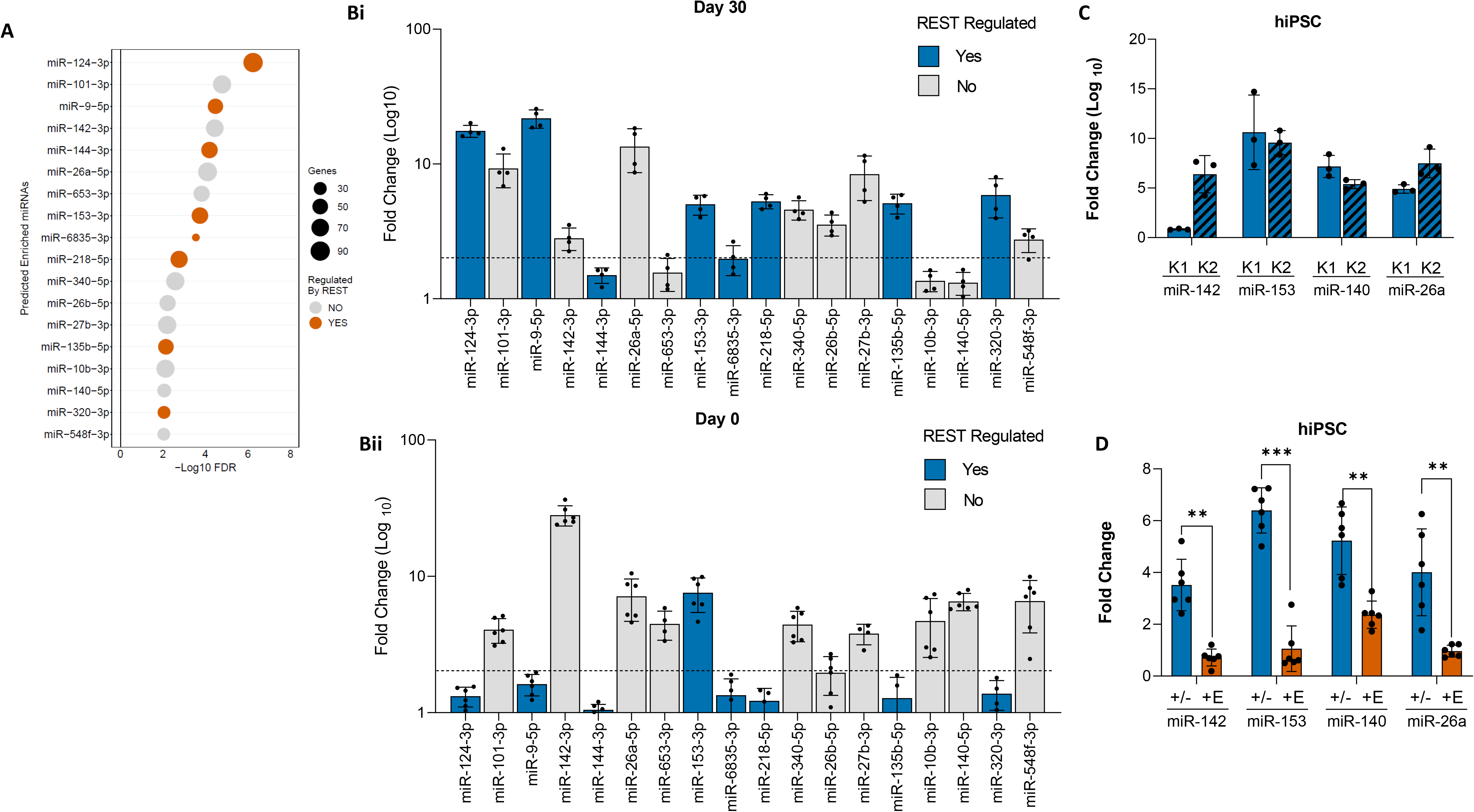
EHMT1 regulates REST expression through a specific set of miRNAs: **A:** Prediction enrichment analysis of upregulated miRNAs based on EHMT1^+/−^ Day 30 transcriptomic data, where orange miRNAs are those known to be regulated by the REST complex. Prediction was considered significant when P < 0.01, plot size indicates the number of DEGs predicted to be targeted by the relevant miRNA. **B:** Analysis of predicted miRNA expression by qRT-PCR. Mean fold change over vehicle control, n ≥ 4 independent experiments, log10 scale axis. Increased miRNA expression = FC > 3. Day 0 and day 30 hiPSC derived neurons following constitutive treatment with 400nM UNC0638. **(i)** Day 0: Of 18 miRNAs, 11 showed increased expression (61.1%), only one is known to be targeted by the REST complex. **(ii)** Day 30: Of 18 miRNAs, 13 showed increased expression (77.7%), of these 8 are known to be targeted by the REST complex. **C:** Analysis of predicted miRNA expression by qRT-PCR. Mean fold change over vehicle control, n = 3 independent experiments, log10 scale axis. Mature miRNA expression analysis of miRs predicted to target REST by qRT-PCR in patient hiPSCs (K1, K2). Except for miR-142, all miRs were increased in expression in patient lines. **D:** Gene expression analysis by qRT-PCR in EHMT1^+/−^ hiPSCs or EHMT1^+/−^ hiPSCs + exogenous EHMT1 plasmid (+E). Elevated expression of miR-142, miR-153, miR-140 and miR-26a in response to hypomethylation was suppressed by overexpression of EHMT1 protein. Data were presented as Mean ± SEM and analysed by One-way ANOVA with post hoc comparisons using Tukey’s multiple comparisons test comparing to control samples. *P < 0.05, **P < 0.01, ***P < 0.001.

Our predicted miRNA set was validated by qRT-PCR of newly generated Day 0 (pluripotent) and Day 30 (neuronal) RNA samples, which had been treated with 400 nM UNC0638, an EHMT inhibitor that at this concentration causes 50% inhibition of EHMT1 activity (Fig. S3). Thirteen of the 18 predicted miRNAs were confirmed to be upregulated at Day 30 following EHMT1 inhibition (Fig. 2B). The results confirm that reduced EHMT1 activity led to miRNA de-repression during neurodevelopment stages, however, the profile of up-regulated miRNAs differed from the equivalent pluripotent iPSC (Day 0) analysis, with only 8 miRNAs up-regulated at both time points. Strikingly, all 5 miRNA that are unique to the D30 neuronal sample are also REST targets. Only one REST targeted miRNA, miR-153, is expressed at Day 0 and is also up-regulated at Day 30

Among the miRNAs elevated at Day 0, four (miR-142-3p, miR-153-3p, miR-140-5p and miR-26a-5p) were predicted to bind and suppress REST mRNA (Table S2). To confirm that these are specifically linked to EHMT1, we examined their expression in EHMT^+/−^ and KS patient-derived iPSCs, all four miRNAs were upregulated, with the exception of miR-142-3p in patient KS1 (Fig. 2C & 2D). To test whether these miRNAs are directly targeted by EHMT1, we performed ChIP–qPCR for H3K9me2 at their promoters. All showed significant H3K9me2 enrichment, which was lost upon UNC0638 treatment (Fig. S4). Finally, overexpression of the EHMT1 gene was sufficient to reduce the levels of the elevated miRNAs responsible for targeting REST (Fig. 2D).

Based on these findings, we propose that there are two miRNA-mediated regulatory components of REST during neurodevelopment for pluripotent iPSC to neurons. Component 1 comprises a small set of EHMT1-repressed, but non-REST regulated miRNA. These can suppress REST but are not inhibited by REST. Component 2 miRNA both suppress REST and are also repressed by REST; they include miR-9 and miR-124. This Component 2 miRNA are up-regulated later during neurodevelopment at Day 24 and 30. The exception to this pattern miR-153, which has both Component 1 and 2 features, being both inhibited by EHMT1 and REST, and induced by reduced EHMT1 activity in the pluripotent cell state.

### REST protein is regulated by cooperation of multiple miRNAs in pluripotent iPSC

To confirm a direct interaction of our Component 1 miRNA (miR-142-3p, miR-153-3p, miR-140-5p and miR-26a-5p) with REST mRNA, we sought a means to reduce their activity. However, as the target sites of all 4 miRNA do not overlap and it has been previously demonstrated that miRNAs are capable of simultaneously targeting the same mRNA [37], we adopted a strategy based on expression of multimeric miR-sponges [38]. In our approach we transiently expressed a non-coding RNA comprising of a concatemer of antisense sequences to our target miRNA, enabling simultaneous repression of multiple different miRNAs (Fig. S5). Strength of repression for individual miRNAs could be controlled by varying the number of antisense sequences targeting the different miRNA. Each multimiR-sponge was validated by uses of luciferase reporter genes constructed for each miRNA (Fig S5E).

MultimiR-sponges for the four component 1 miRNAs either as a single miRNA type or in combinations of three different miRNAs were introduced into UNC0638 treated-iPSC at the pluripotent stage (Day 0) and expression of the RE1-repressed target genes NRXN3, ACTA1 and CALB1 was measured (Fig. 3A-C). Suppression of the miRNA would be expected to block UNC0638-induced expression of each mRNA for all REST target genes. Strong suppression was observed in the presence of any multimiR-sponge that combined miR-153-3p with any other pair-wise combination of the other three miRNAs. The three-miRNA multimeric combination of miR-142-3p, miR-140-5p and miR-26a-5p in the absence of miR-153-3p or singularly targeting miR-153-3p alone failed to deliver statistically significant suppression. These observations indicate that strong suppression of REST genes requires the concerted action of miR-153-3p and two other miRNAs. Strongest suppression of RE1-containing target genes was seen with the combined suppression of miR-140, miR-26a and miR-153, with relative FC of 1.56±0.56, 1.10±0.30 and 2.37±0.42 for NRXN3, ACTA1 and CALB1 respectively. Targeting of REST protein was confirmed by Western blot analysis of UNC0638 treated-hiPSC^+/+^ or EHMT1^+/−^ mutant cells. In both cases, a decrease of REST protein was reversed in the presence by expression of the 140/26a/153-multimiR sponge (Fig. 3D & 3E). Collectively, these data indicate that down-regulation of REST protein in pluripotent iPSCs is mediated by the cooperative action of a small set of miRNAs.

**Figure 3:**
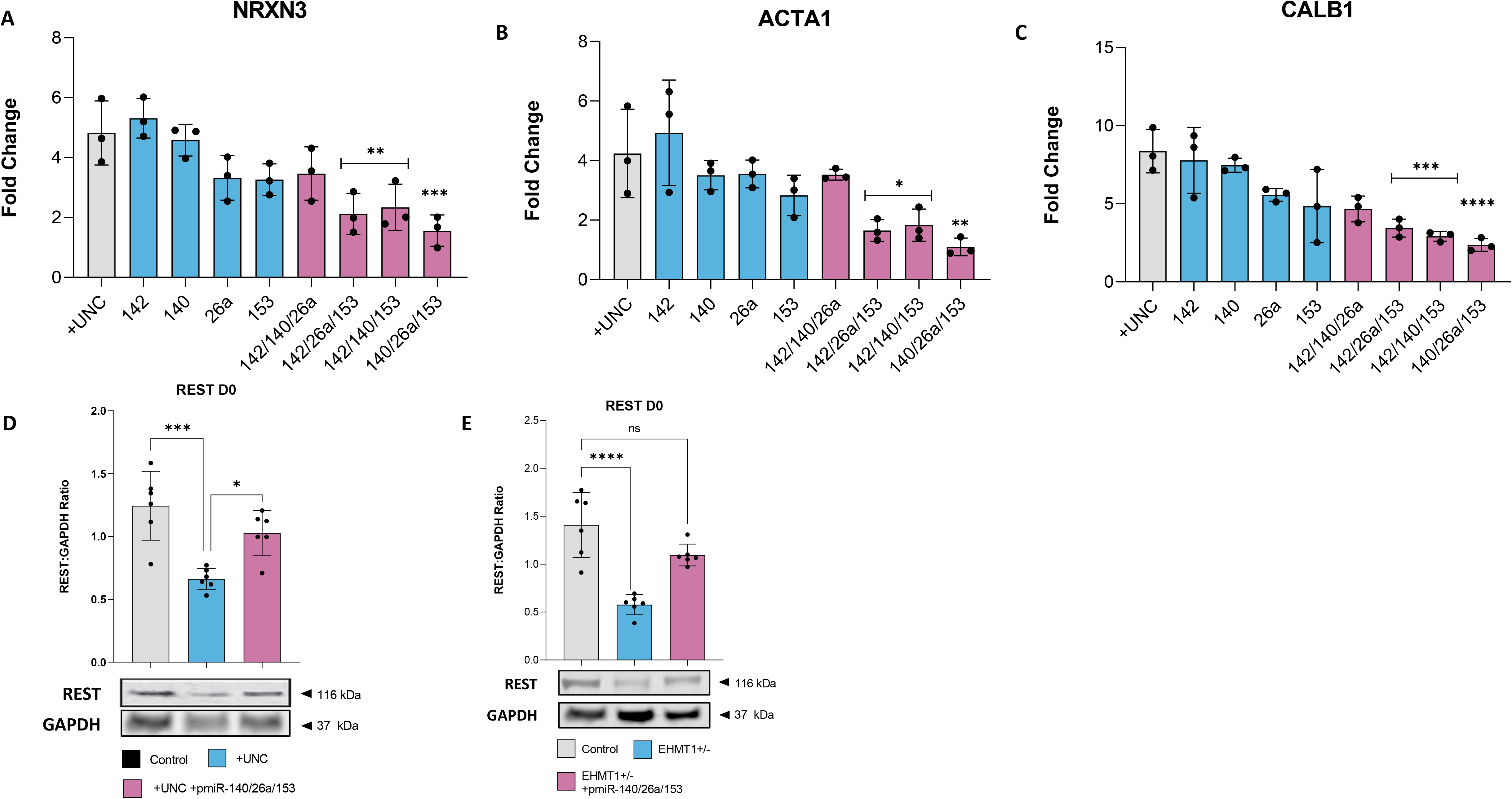
REST is cooperatively regulated by miRNAs: **A-C:** Analysis of REST targets by qRT-PCR in hiPSCs treated with 400nM UNC0638 and varying miR-sponge combinations, as compared to empty-vector (EV) sponge control. Mean fold change over vehicle control, n = 3 independent experiments, log10 scale axis. Statistics: one-way ANOVA across sponge groups within each target gene, with Dunnett’s multiple comparisons versus EV control. **A:** NRXN3 expression, only multimiR-sponges containing miR-153 suppressed increased expression. **B:** ACTA1 expression, only multimiR-sponges containing miR-153 suppressed increased expression. **C:** CALB1 expression, all multimiR-sponges could reduce increased target expression, in addition to miR-153 alone. n ≥ 3 independent experiments, log_10_ scale axis. **D:** REST protein in control hiPSCs treated with UNC0638 (4 days) and infected with EV sponge versus multimiR-140/26a/153 sponge. Normalised to GAPDH. n = 6. Statistics: two-tailed unpaired Welch’s t-test (EV vs multimiR). **E:** REST protein in EHMT1^+/−^ hiPSCs versus isogenic control hiPSCs, with EV or multimiR-140/26a/153 sponge as indicated. Normalised to GAPDH. n = 6. Statistics: two-way ANOVA (genotype × sponge) with Šidák multiple comparisons for the planned contrasts (e.g., EV vs multimiR within each genotype; EHMT1^+/−^ vs control within each sponge condition).

### Increased miR-9 and miR-124 expression accelerates neuronal differentiation

miR-9 and miR-124 are two prominent Component 2 miRNAs that are well established as regulators of neuronal development [35, 39]. To examine their regulation in the context of EHMT1 activity, we followed their abundance over 70 days of neuronal differentiation, comparing EHMT1^+/−^ mutant iPSC to control cells (Fig. 4A-B). In contrast to our Component 1 miRNAs, neither miR-9 nor miR-124 showed a significant increase in the EHMT1^+/−^ mutant iPSC during the first 10 days of differentiation. However, by Day 20 we observed the emergence of a significant, multi-fold increase of both miR-9 and miR-124. This higher expression level persisted until Day 60, at which point miRNA expression in the control cells began to approach those of the EHMT1^+/−^ mutant.

**Figure 4:**
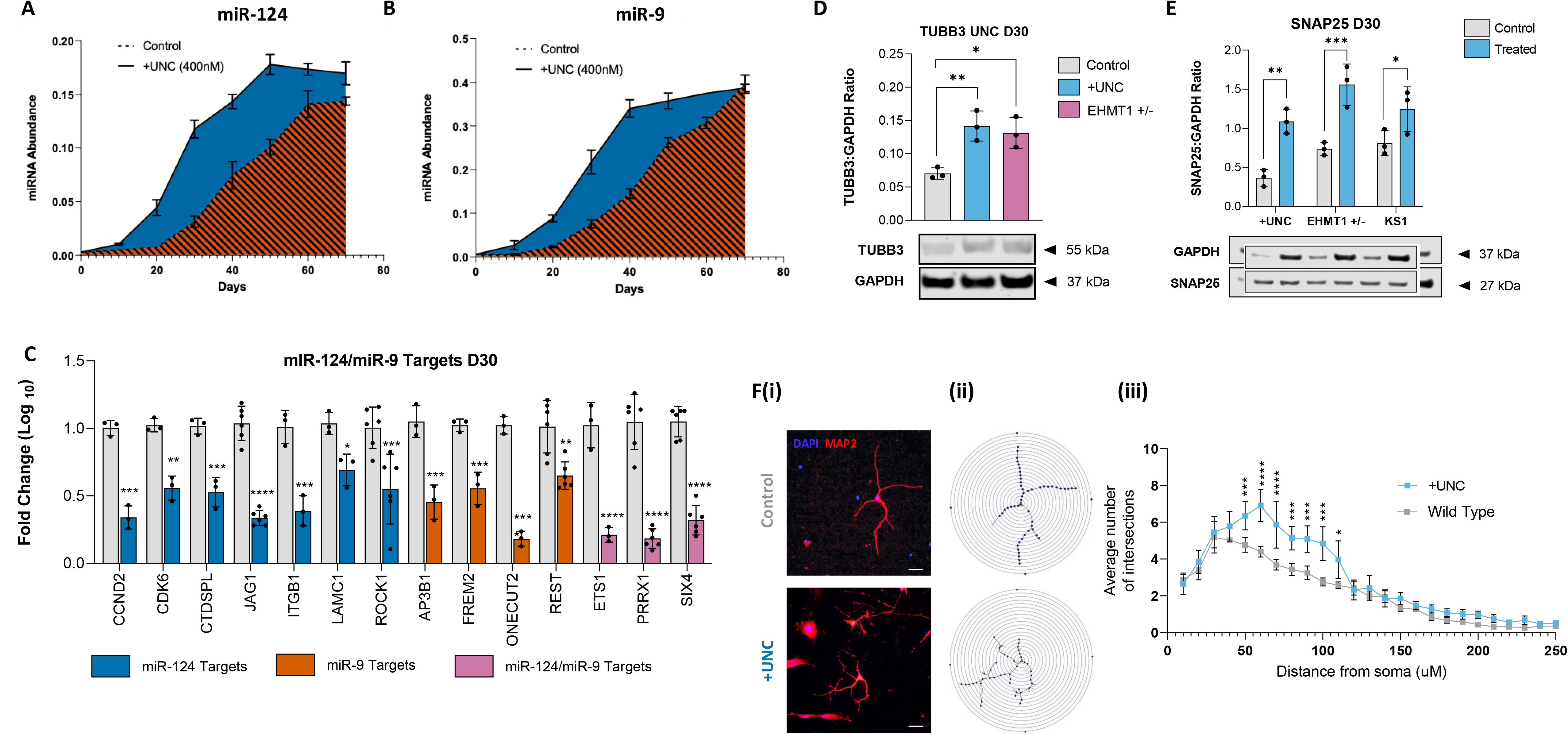
REST dysregulation impacts brain miRNAs and neuronal development: **A,B:** Relative miRNA abundance miR-9 and miR-124 in EHMT1^+/−^ hiPSC derived neurons compared to controls. Samples from Day 0-70 were measured by qRT-PCR, n ≥ 3 independent experiments. **C:** Analysis of miR-124 and miR-9 validated mRNA targets by qRT-PCR in EHMT1^+/−^ hiPSC derived neurons at day 30. Targets are all decreased in comparison to control cells (n ≥ 3 independent experiments). **D:** Western blot analysis of TUBB3 protein expression in hiPSCs treated with UNC0638 and EHMT1^+/−^ hiPSCs, differentiated to neurons (Day 30) in comparison to vehicle control or isogeneic control cells respectively. TUBB3 expression was significantly increased in cells with reduced EHMT1 activity. **E:** Western blot analysis of pre-synaptic SNAP25 protein expression in hiPSC treated with UNC0638, EHMT1^+/−^ hiPSCs and patient KS hiPSCs, all differentiated to Day 50 neurons and relative to vehicle control or isogenic controls. All three showed significant increases in SNAP25 expression. **F:** (i) Representative immunofluorescence images of hiPSC-derived neurons stained for MAP2 (red) and DAPI (blue) in control and UNC0638-treated (+UNC) conditions at day 30, illustrating neuronal morphology and neurite outgrowth. Scale Bar, 70 μm. (ii) Corresponding neuronal reconstructions and branching skeletons generated from MAP2-positive cells, used for Sholl analysis to quantify neurite complexity. (iii) Quantitation of neurite branching complexity by Sholl analysis, shown as the average number of intersections as a function of distance from the soma (µm). UNC0638-treated neurons exhibit increased neurite complexity compared to wild-type controls. Data were presented as Mean ± SEM from n = 3 independent experiments. Statistical analysis was performed using a two-way repeated measures ANOVA (factors: treatment × distance from soma) with appropriate post hoc multiple comparisons testing, with significance indicated (*p < 0.05, **p < 0.01, ***p < 0.001).

To confirm that these gene regulatory changes impact on neurodevelopment, we investigated 14 validated mRNA targets of the miR-9 or miR-124, assessing their expression at day 30 in EHMT1^+/−^ derived neurons (Fig. 4C). These included REST protein itself and CTDSPL, a component of the REST complex [35], as well as targets that are typically downregulated in mature neurons, such as ONECUT2 and SIX4 that prevent premature differentiation [40] and JAG1 that controls the balance between NPCs and mature neurons [41]. All 14 target mRNAs were decreased in EHMT1^+/−^ derived Day 30 neurons. At the protein level, the pan-neuronal marker TUBB3 was significantly increased at Day 30 in UNC0638-treated and EHMT1^+/−^ neurons (Fig. 4D) and the pre-synaptic marker SNAP25 was significantly increased at Day 40 in UNC0638-treated, EHMT1^+/−^ and patient-derived neurons (Fig. 4E). At the cell level, quantitative Sholl analysis at Day 40 indicated an increase in branching complexity in response to EHMT1 depletion, with UNC0638-treated treated cells showing significantly greater number of average intersections (Fig. 4F).

Taken together, these results indicate that reduced EHMT1 activity is associated with up-regulation of neuronal-specific miRNA, miR-9 and miR-124 and suppression of REST-targeted brain miRNAs and precocious neuronal gene expression. This commences at the transition from NPC to neurons and persists throughout neurodevelopment.

### Component 1 miRNA are both necessary and sufficient switch to trigger neurodevelopment of human iPSC

A simple hypothesis is that up-regulation of Component 1 miRNA initiates neuronal differentiation, which is then locked into place by up-regulation of Component 2 miRNAs. To explore the dependency between Component 1 (miR-142-3p, miR-153-3p, miR-140-5p and miR-26a-5p) and Component 2 (miR-9 and miR-124) we generated a retroviral multimiR sponge to deliver a stable and persistent inhibition of miR-153-3p, miR-140-5p and miR-26a-5p (Fig. 5A). Following continuous UNC0638-treatment we observed repression of REST protein in control cells that express a scrambled sequence miRNA, but this was fully blocked in neurons that express a 153/140/26a-5p multimiR sponge (Fig. 5B). Consistent with the requirement for an increase in component 1 miRNA to trigger Component 2 miRNA, UNC0638-induced increase of miR-9 or miR-124 was blocked in a 153/140/26a-5p multimiR sponge (Fig. 5C). Finally, protein expression of NEFM and SYN1, two REST repressed genes, was elevated in UN0638-treated control cells, but again this was blocked by expression of a 153/140/26a-5p multimiR sponge (Fig. 5D,E).

**Figure 5:**
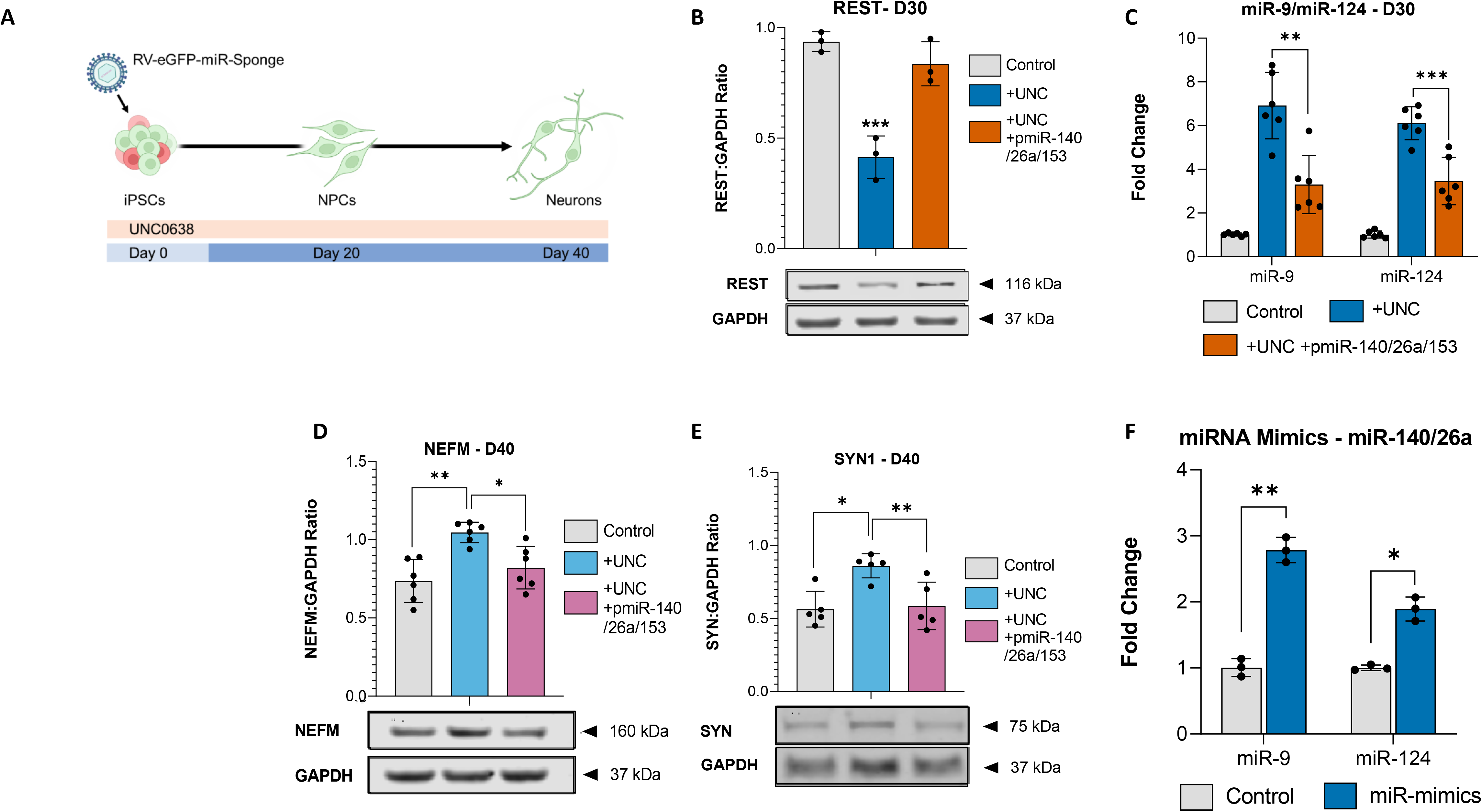
Elevated miR-142, miR-153, miR-140 and miR-26a miRNA are necessary and sufficient for neuronal differentiation: **A:** Schematic showing experimental design for stable suppression of EHMT1-miRNAs following infection with retroviral multimiR sponge. **B:** Western blot analysis of REST protein expression in hiPSC derived neurons (day 30) treated with 400nM UNC0638 (UNC), samples infected with retroviral multimiR-sponge against miR-140/26a/153, and compared relative to vehicle only controls. Treatment with retroviral multimiR-sponge returned REST protein levels to normal. **C:** miRNA expression analysis of miR-9 and miR-124 by qRT-PCR in hiPSC derived neurons at day 30 following treatment with UNC0638 and samples infected with retroviral multimiR-sponge against miR-140/26a/153, as relative to vehicle control. n = 6 independent experiments. **D, E:** Western blot analysis of neuronal proteins NEFM (D) and SYN (E) in hiPSC derived neurons (Day 40) infected with retroviral multimiR-sponge against miR-140/26a/153 and treated with 400nM UNC0638. **F:** miRNA expression analysis of miR-9 and miR-124 by qRT-PCR in hiPSC derived Day 30 neurons following introduction of miRNA mimics of miR-140 and miR-26a, relative to scramble a sequence control (n = 3 independent experiments). Data were presented as Mean ± SEM and analysed by one-way ANOVA with post hoc comparisons using Tukey’s multiple comparisons test comparing to control samples. *P < 0.05, **P < 0.01, ***P < 0.001.

These observations show that in combination Component 1 miRNA are necessary for miR-9 and miR-124 regulation of REST function. To establish whether they are sufficient, we introduced synthetic miRNA-mimics for miR-26a and miR-140 into Day20 NPC cells and measured miR-9 and miR-124 expression in Day 30 neurons. miR-153 was not used in this experiment due to its regulation by REST, which would confound the experiment. Transfection of control NPC cultures with synthetic miRNA-mimics for miR-26a and miR-140 led to a significant increase of both miR-9 and miR-124 (Fig. 5F). These data demonstrate that introduction of exogenous miR-140-5p and miR-26a-5p, likely by cooperating with endogenous miR-153-3p, is sufficient to drive miR-9 and miR-124 expression and would be expected to suppress REST and de-repress neurodevelopment.

Overall, these results indicate that elevation of Component 1 miRNAs (miR-142-3p, miR-153-3p, miR-140-5p, miR-26a-5p) is both necessary and sufficient to reduce REST protein expression to promote progress into neuron cell states. In EHMT1^+/−^ hemizygous engineered cells; wild type cells following UNC0638-treatment or KS-patient cells, reduced EHMT1 activity elevates Component 1 miRNA expression. This partially lowers REST, leading to early and higher levels of Component 2 miRNA, which in turn result in premature neuronal differentiation (Fig. 6).

**Figure 6:**
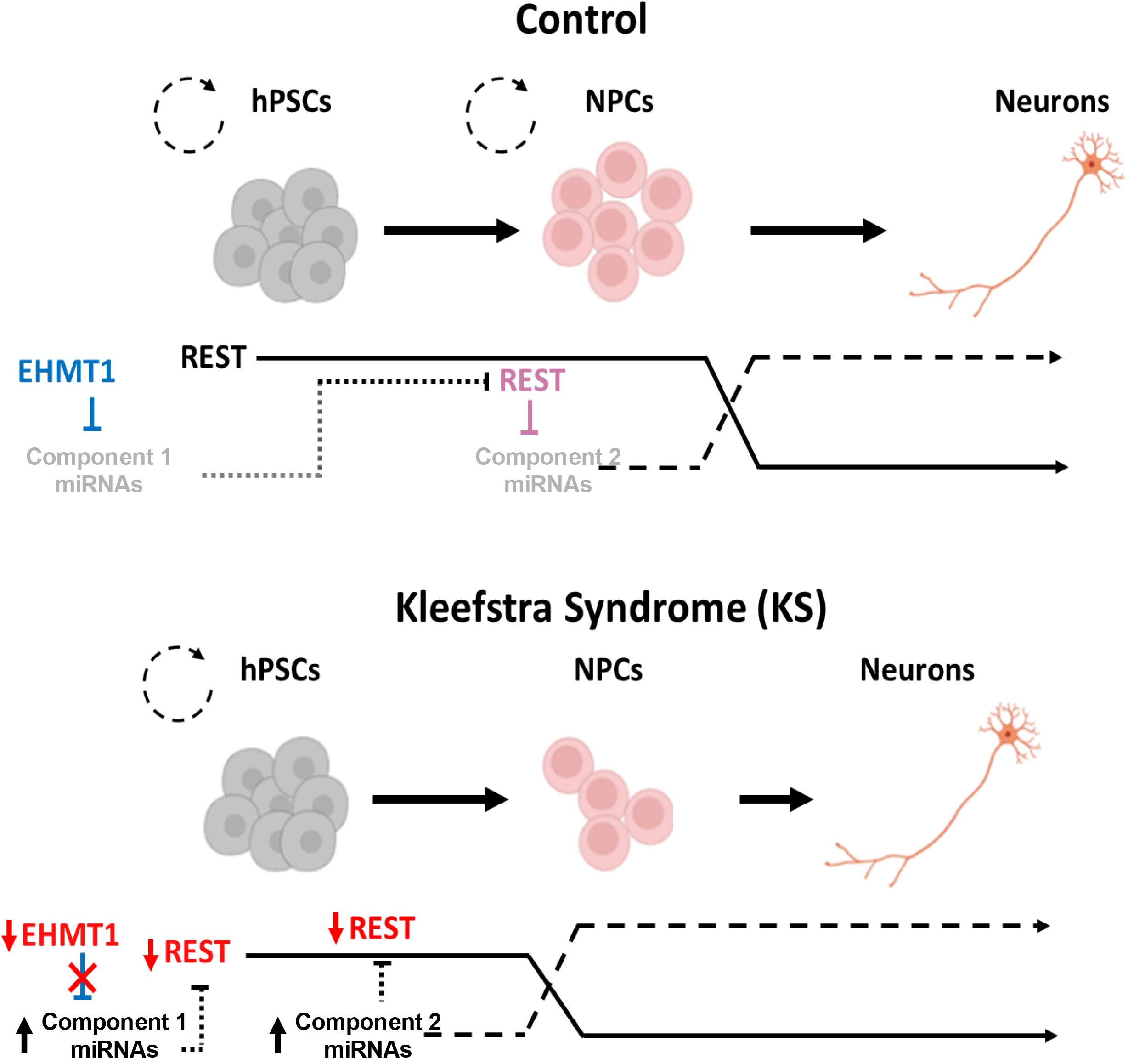
Scheme to describe how altered EHMT1 activity leads to premature neuronal differentiation. In cells with normal levels of EHMT1 activity (designated as Control) Component 1 miRNAs miR-142-3p, miR-153-3p, miR-140-5p and miR-26a-5p) are initially suppressed by EHMT1 mediated H3K9 dimethylation, maintaining high levels of REST protein, blocking the transition from NPC to developing neurons. When component 1 miRNA reach a threshold level in NPC the begin to reduce REST protein levels to a point where Component 2 miRNA (eg miR-9 and miR-124) also begin to be expressed. This leads to a rapid decrease in REST protein and a switch to a neurodevelopmentally permissive state. In cells where EHMT1 is reduced, such as in KS, REST protein is permanently at a lower level, and cells reach the threshold to increase

### Early miRNA expression confers directionality to neuronal differentiation

A key remaining question is why neuronal differentiation proceeds via a complex two-component process? We propose that this mechanism makes the transition from pluripotent cells to neuronal differentiation irreversible, imparting directionality to neurodevelopment. To test this idea, we transiently treated cells with UNC0638, from Day 0 to Day 10 (early NPCs) and then withdrew the inhibitor (Fig. 7A). Western analysis showed that following inhibitor withdrawal there was an overall recovery of H3K9me2 (Fig. S6), but no restoration of control of levels of REST protein, which remained decreased, or of miR-9 and miR-124, which remained elevated, (Fig. 7B & C). This indicates that once the mechanism has been triggered it continues, even in the absence of the molecular initiating event.

**Figure 7:**
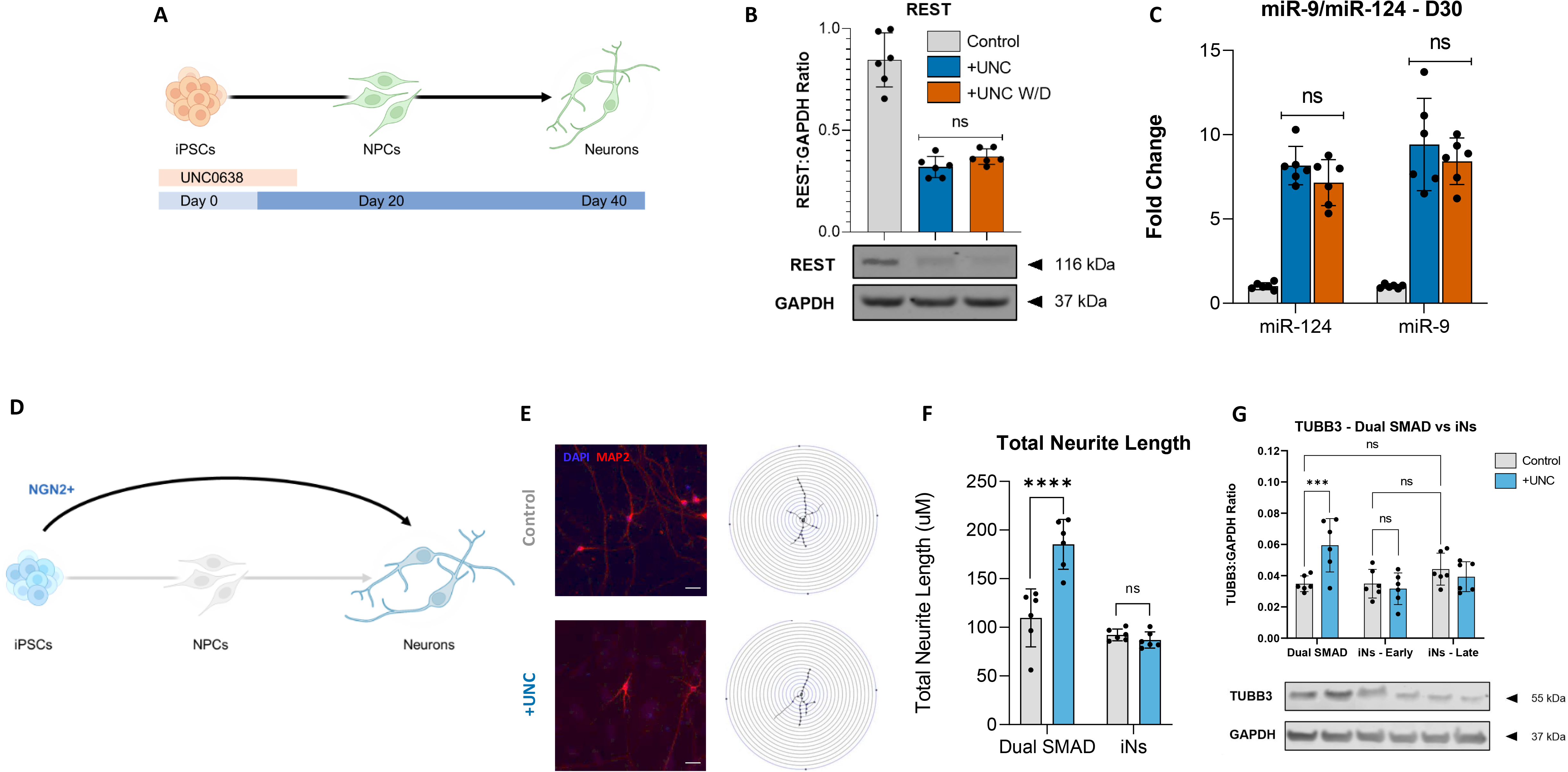
miRNA control of neurodevelopment via NRSF/REST regulation: **A:** Schematic showing experimental design for (i) UNC0638 withdrawal at day 10 and **B:** Western blot analysis of REST protein expression in hiPSC derived neurons at day 30 following treatment with UNC0638 (UNC) from days 0-10, as compared to vehicle control. Withdrawal of UNC0638 did not restore REST expression. **C:** Mature miRNA expression analysis of miR-9 and miR-124 by qRT-PCR in hiPSC derived neurons at day 30 following treatment with UNC0638 from days 0-10, as compared to vehicle control. n = 6 independent experiments. Withdrawal of UNC0638 did not reduce miR-9 or miR-124 expression. **D:** Induction of iPSCs to maturing neurons by forced expression of an inducible NGN2 transgene, bypassing NPC stages. **E:** Representative immunofluorescence images of induced neurons stained for MAP2 (red) and DAPI (blue) in control and UNC0638-treated (+UNC) conditions at 20 days post induction (DPI). Scale Bar, 70 μm. Corresponding neuronal reconstructions and branching skeletons generated from MAP2-positive cells, used for Sholl analysis to quantify total neurite length. **F:** Quantification of total neurite length (µm) in neurons generated either by Dual SMAD differentiation (day 30) or NGN2-induced direct conversion (15 days post-induction), treated with vehicle control (grey) or UNC0638 (blue). UNC0638 treatment significantly increased total neurite length in Dual SMAD-derived neurons, while no significant effect was observed in NGN2-induced neurons (iNs). **G:** Western blot analysis of TUBB3 protein expression (normalised to GAPDH) in neurons generated by Dual SMAD differentiation (day 30), early NGN2-induced neurons (iNs; day 15 post-induction), and late NGN2-induced neurons (iNs; day 30 post-induction), treated with vehicle control (grey) or UNC0638 (blue). UNC0638 treatment significantly increased TUBB3 expression in Dual SMAD-derived neurons, while no significant differences were observed in early or late iNs. Data are presented as mean ± SEM from n = 6 independent experiments.

Finally, to confirm that this is indeed a true developmental “switch” that controls progression of neurodevelopment rather than an additive effect overlaying different neurodevelopmental processes. We bypassed the NPC stage by driving immediate neuron formation using NGN2 expression to generate induced neurons (iNs) and then comparing outcome in the presence or absence of EHMT1 inhibition to those of cells developed using the conventional (dual SMAD method) differentiation induction, which proceeds via the initial generation of NPC (Fig. 7D). In dual SMAD cultures, UNC0638 treatment resulted in increased neurite length and elevated TUBB3 expression compared to control (Fig. 7E & F). In contrast, UNC0638-treated iNs showed no significant differences in neurite complexity, total neurite length, or TUBB3 protein levels compared to controls (Fig.7G, Fig. S7).

Collectively, these data indicate that the switch operates within early neurodevelopmental stages, acting as regulator of the timing of neuronal differentiation. The further consequence of its two-component configuration is to make the switch from neurodevelopmental repression to a permissive state irreversible, conferring directionality to neurodevelopment.

## Discussion

In this study we define an EHMT1–miRNA–REST regulatory network that creates an irreversible developmental switch and imparts directionality to *in vitro* neurodifferentiation of iPSC. The molecular basis for this regulatory mechanism derives from the interaction of miRNA with two epigenetic repressors, EHMT1, which is responsible for the H3K9me2 chromatin mark, and REST, which represses neuronal gene expression and neuronal differentiation[20]. To explain the operational principles of the network, we propose that there are three key mechanistic components to the miRNA-mediated switch: (i) a double negative Feedback loop (FBL), centred on miRNA-REST interactions; (ii) an overlapping Feedforward loop (FFL) [43]; and (iii) an initiating step, in this case reduced EHMT1 activity (Fig 8). These are discussed in more detail below.

**Figure 8:**
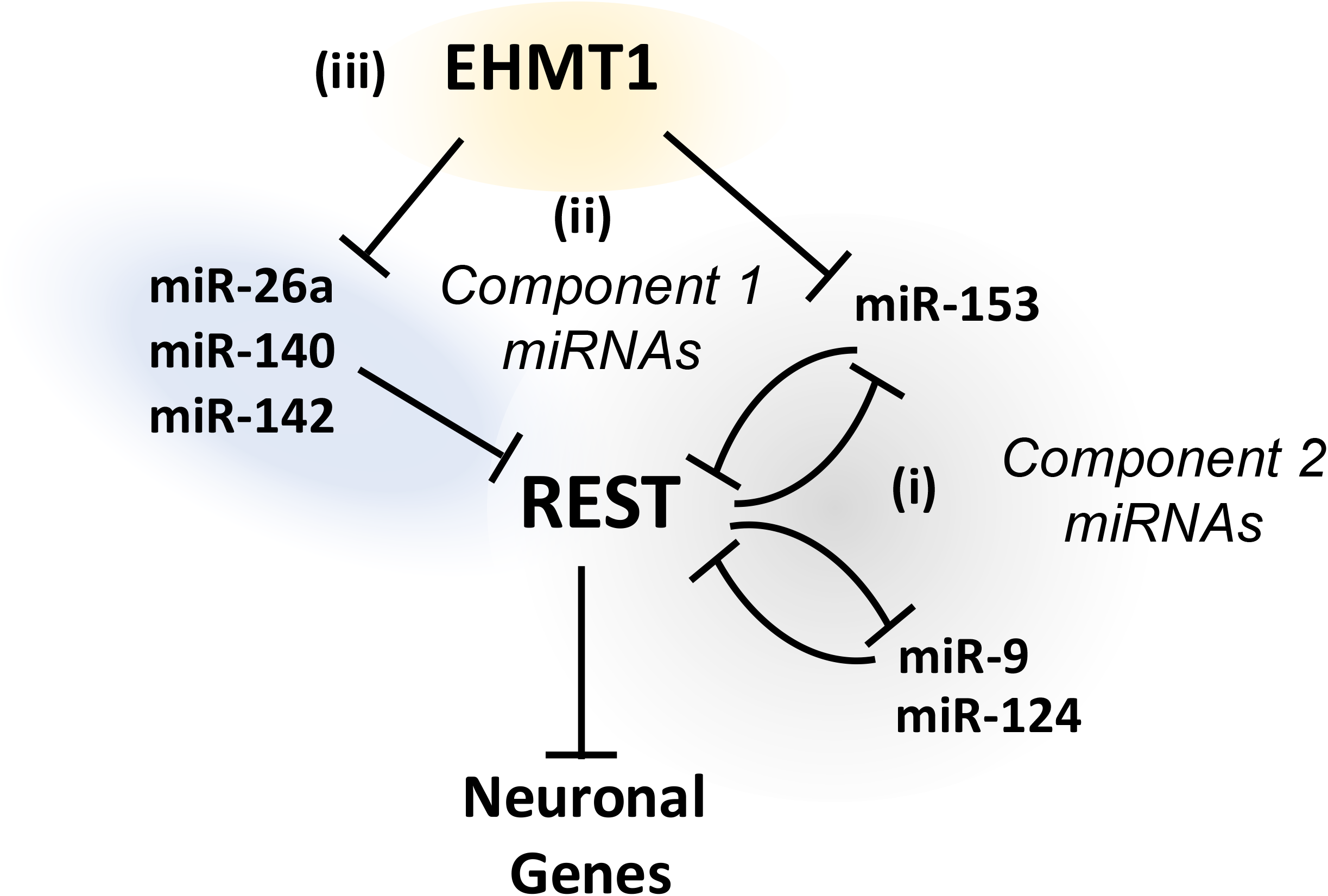
Schematic illustration of the miRNA interactions that control the switch from repressive to permissive neurodevelopmental states. REST protein represses neuronal gene transcription to block neurodevelopment. REST levels can be controlled through the action of both (i) a negative Feedback Loop (FBL) via Component 2 miRNA and (ii) a Feedforward Loop (FFL) via Component 1 miRNA. In combination, these two control loops create a unidirectional switch from a high REST repressed state to a low REST permissive state. The timing of the switch is dependent on epigenetic repression by EHMT1 (iii), whose loss leads to premature neuronal maturation.

### A negative feedback loop controlling neurodifferentiation

Central to the genetic switch is its double negative FBL, in which miRNAs miR-9 and miR-124 are inhibited through the action of the REST complex, but in turn target REST or Co-REST mRNA, blocking protein translation or degrading REST mRNA [35, 39, 44, 45]. In this configuration, high levels of REST protein will suppress miR-9 and miR-124 thus stabilising REST-mediated gene repression, whereas in low levels of REST, miR-9 and miR-124 expression is high, and the REST-complex is maintained at a low level. This creates two states, one repressive and one permissive for neuronal differentiation (Fig 8i).

In support of this mechanism, both miR-9 and miR-124 are strong repressors of REST and CoREST respectively and hence are potent drivers of neuronal differentiation. In this report we observed that mature markers of neuronal differentiation, MAP2, TUBB3 and SNAP25 were prematurely increased when REST protein was lower. Accelerated neuronal differentiation is also seen in cells when REST is knocked down or when miR-9 and miR-124 are exogenously expressed [39]. In the mouse brain, premature overexpression of miR-124 alone leads to the reduced expression of the laminin receptor genes laminin γ1 (LAMC1) and integrin β1 (ITGB1), culminating in structural abnormalities [46]. In human iPSC we observed miR-124 associated decreases of LAMC1 and ITGB1 gene expression, alongside decreased expression in a range of cell cycle regulators, such as CCND2, CDK6 and JAG1. Importantly these targets are required to maintain the balance between neuronal progenitor populations and mature neurons [41, 47]. Our findings are consistent with the proposed roles of miR-124 and miR-9 as a regulator of neurodifferentiation.

### An overlapping Feedforward loop

A major feature of double negative FBLs is that they form and maintain robust network states. Although there is plenty of published evidence for robust repression of neuronal gene expression in non-neuronal cell types and neurodifferentiation following REST repression, the question is what mechanisms control the switch from one state to another? We propose this is mediated by a overlap with an FFL driven by increased expression of the Component 1 miRNA.

Here, we show that when expression of our four Component 1 four miRNAs increases, they work in combination to repress REST protein expression and trigger *in vitro* neurodifferentiation. A key element to this regulation is that three of the four miRNAs do not possess an RE1 site and hence are not subject to REST-mediated gene repression. Such a configuration establishes a coherent FFL that confers the properties required to drive the REST-mediated bistable switch (Fig. 8ii). This FFL is asynchronous due to possession of both direct and indirect (via its interaction with FBL) arms and can only cause a change of state when both combine to reach a required threshold. Below threshold, the network is configured to buffer against small circuit fluctuations arising from system noise. This latter feature is further embedded in the FFL due to a need for cooperation of multiple, moderately active miRNA inhibitors, requiring two miRNAs from the set of miR-26a, miR-140, miR-142 and the presence of miR-153. In this respect we note the unusual positioning of miR-153 [48], which is repressed both directly by a H3K9me2 and indirectly by REST.

The full significance of this dual regulation of miR-153 is not clear, but we suggest that it may act to accelerate the switch between repressive and permissive neurodevelopmental states. In this context, our *in vitro* experimental observations show that miR-9 and miR-124 only begin to rise after Day 10 of neurodevelopment, yet we know that increasing Component 1 miRNA expression in the pluripotent state is sufficient to reduce REST protein and elevate expression of some mRNAs normally subjected to REST repression. This difference in timing may reflect the additional time taken for miR-153 to first be derepressed due to lower H3K9me2 and then a subsequent need for miR-153 expression to increase further via lowering its level of REST-mediated suppression. This additional complexity due to interaction of FBL and FFL may explain how timing of neurodevelopment is controlled and once triggered can only proceed in one direction (Fig 7A-C).

### Initiation via loss of EHMT1 activity

The final component of the switch is an initiator, an external input that ultimately leads the pluripotent cells to progress into neurodevelopment. Here, our initiator is the reduction of EHMT1 activity and subsequently the strongly repressive chromatin mark, H3K9me2/3 (Fig 8iii). This can be achieved either acutely using the chemical inhibitor, UNC0638 or chronically via loss of one gene copy. In untreated or non-mutant cells EHMT1 expression is relatively constant from the pluripotent cells to the end of the pluripotent state and then begins to decline during neuronal maturation [3]. Although this expression profile does not exclude developmental regulation of H3K9me2 at a small number of specific gene loci, it argues against a global regulation of EHMT1 as the initiator of neurodevelopment under normal circumstances. In the experiments we describe here, loss of EHMT1 may cause its effects by simply lowering the increase in Component 1 miRNA expression needed to reach the threshold at which the FBL within the switch is triggered (Fig 6).

### Significance for neurodevelopment and NDD

Work has previously demonstrated miRNAs such as the miR-26 family sit upstream of the REST protein, where their knockout impacts both REST-controlled miRNAs and neuronal maturation [49]. However, to the best of our knowledge, this report is the first to show the need for additional miRNAs to act cooperatively during regulation of REST expression. Similar endogenous cooperative miRNA binding has been demonstrated in HEK cells, where the combined presence of two miRNAs, but not their individual counterparts, is required for gene repression [50]. Likewise cooperative miRNA repression has been shown to be indispensable for *in vivo* silencing within C. elegans [51]. This synergistic binding likely enables fine control of protein output in an energy-efficient manner, ensuring adaptability in a robust model. In this manner, EHMT1 repression acts to prolong the initial plastic NPC state, an essential step in the highly protracted process of human neurodevelopment [52].

Conversely, this extended developmental period may increase susceptibility to neurodevelopmental disorders, as substantiated by the dysregulation of various miRNAs in NDDs. Our EHMT1 regulated miRNAs are not limited to KS; miR-140 and miR-142 have been found to be elevated in patients with Autism [53, 54], miR-26 has routinely shown to be involved with Schizophrenia patients [55, 56] and miR-153 has been associated with epilepsy and Autism [57, 58]. The role of EHMT1 and its associated miRNAs in neurodevelopmental disorders underscores the delicate balance between epigenetic factors and miRNA networks in shaping the timing and progression of neural differentiation, highlighting the vulnerability of the human brain to disruptions in this process.

**In summary**, this study has identified an EHMT1-sensitive, miRNA network, that is responsible for the disruption of a REST-miRNA genetic switch. The work highlights the delicate reciprocal balance between epigenetic regulators, miRNAs and their emergent properties via Feedback and Feedforward loops to control the timing of neuronal development. Importantly, it highlights the role of EHMT1 in regulating the progression of neurogenic development.

## Materials and Methods

### Human iPSC culture and neuronal differentiation

The IBJ4 human iPSC line derived from the BJ fibroblast cell line (ATCC: CRL-2522) was used, unless indicated. HiPSCs were grown on cultrex (ThermoFisher Scientific) in Essential 8^TM^ medium (ThermoFisher scientific) at 37°C, 5% CO_2_. Medium was changed every day, and cells were passaged using gentle cell dissociation reagent (Stemcell Technologies). Differentiation of hiPSCs to glutamatergic neurons was based on a modified method of Chambers et al. [59], using N2B27 (2/3 DMEM/F12; 1/3; neurobasal; B27-RA; N2; 1xPSG; 0.1 mM β-mecaptoethanol) + 100 nM SB431542 and 100 nM LDN193189, followed by a N2B27 with B27 + retinoic acid (with 10 µM DAPT for first 7 days) on PDL (Sigma)/Laminin (Roch) at 200000 cells/cm^2^.

### Expression Analysis

*qRT-PCR*: Total RNA was extracted from cells using the miRNeasy mini kit (217004, Qiagen, Germany). For each miRNA sample, 200 ng of total RNA was reverse transcribed using the miRCURY LNA RT Kit (339340, Qiagen, Germany), whilst for each mRNA sample 1 µg of total RNA was reverse transcribed using the High-Capacity RT cDNA Kit (ThermoFisher Scientific). qRT-PCR analysis of miRNAs used the miRCURY LNA SYBR green kit (Qiagen, Germany) and mRNA analysis used the qPCRBIO SyGreen Blue Mix (PCR Biosystems, UK). All qRT-PCR reactions were performed in triplicate on a StepOnePlus^TM^ Real-Time PCR System (Applied Biosystems) and relative expression calculated using the 2^−ΔΔCT^ method with data normalized to GAPDH or SNORD48 (see Table S1 for primer sequences).

*Western blot analysis*: Cells were lysed in RIPA buffer (Sigma) and protease inhibitor cocktail for 30 mins at 4°C. Cell supernatants were collected, before LDS sample buffer (ThermoFisher Scientific) and sample reducing agent (NuPAGE) were added and samples were heated to 95°C for 5 mins. A total of 20 µg of protein per sample was separated by electrophoresis on 4-12% Bis-Tris Plus Gels (Life Technologies), transferred to nitrocellulose, blocked with a milk powder solution, 5% (w/v) in Tris-buffered saline containing 0.1% Tween-20 (v/v) (TBST) for 60 mins at RT and incubated overnight with primary antibody against REST (1:500) (07-579, Merck), H3K9me2 (1:500) (17-648, Merck), diluted in blocking solution. After washing in TBST blots were incubated with an appropriate IRDye®-conjugated secondary antibody (Li-COR) and visualized/quantified with a Licor/Odessey infrared imaging system (Biosciences, Biotechnology). All data normalization was against GAPDH.

### Bioinformatic Analysis

Microarray and RNA sequencing datasets of gene expression studies focussed on *EHMT1* depletion or mutation were accessed and downloaded from Gene Expression Omnibus (GEO) Datasets available on the National Centre for Biotechnology Information (NCBI) Database website (http://www.ncbi.nlm.nih.gov/gds/). The search words used were “*EHMT1*” and “Kleefstra Syndrome” and through filtering the search was restricted to *Homo Sapiens*. From this search the raw data for a single dataset was downloaded (GSE178646, n=3).

Counts were read into R 4.3.2 (https://www.R-project.org/) and filtered to retain genes with counts greater than 10. Principle Component Analysis Plots indicated no outlying samples. Differential gene expression (DGE) analysis was performed using DESeq2 (R package, v1.30.0), where the difference in gene counts was assessed between the wild type control and SNP samples. P values were adjusted for multiple comparisons using Benjamini-Hochberg correction address false discovery rate (FDR). A gene was considered differentially expressed if it has an adjusted P-value of less than 0.05 and a Log2 fold change of ±1.5 or greater.

#### Functional Enrichment Analysis

For enrichment analysis, significant genes from RNA seq analysis were grouped into either “upregulated” and “downregulated” datasets. Enrichment analysis was performed for each gene set to determine Gene Ontology – Biological Processes (GOBP), using the fgsea multilevel enrichment test and genes were ranked by minimum significant difference signed msd (fgsea). Enriched terms with a padj value less than 0.01 were considered significant. Enriched GOBP terms were clustered based on semantic similarity using GOSemSim (R package, v2.24.0), and clusters were manually summarized. Disease-gene network analysis was performed using the DisGeNET database (V7.0) [60], calculating overlap significance for DEGs and disease genesets by hypergeometric probability, with adjusted P<0.001 considered statistically significant. Tissue enrichment analysis was conducted using the TissueEnrich (R package, v1.20.0), with enrichment of tissue specific gene sets assessed within KS DEGs.

#### Definition of RE1/REST gene list

Genes annotated as RE1/REST-associated were derived from public resources by integrating ENCODE REST (NRSF) ChIP-seq annotations with ChEA “Consensus TFs from ChIP-X” REST/NRSF target gene sets. Where required, REST binding sites were assigned to genes using a nearest-transcription-start-site rule (nearest TSS) / [within 10 Kb], and gene identifiers were standardised to Ensembl IDs. Duplicates were removed and the union of genes supported by either resource was used as the RE1/REST gene list for downstream enrichment testing.

#### RE1 enrichment permutation test

To test whether RE1/REST-associated genes were enriched among upregulated DEGs beyond chance expectation, we performed a permutation-based overlap analysis. Upregulated DEGs were defined as genes with DESeq2 FDR-adjusted P < 0.05 and log2FC ≥ 1.5. The background “gene universe” comprised genes expressed in the dataset (baseMean > 10). We computed the observed number of RE1/REST genes among upregulated DEGs and compared this with a null distribution generated from 10,000 random gene sets of equal size sampled without replacement from the expressed-gene universe. The empirical P value was calculated as (r + 1)/(n + 1), where r is the number of permutations with overlap ≥ the observed overlap and n is the number of permutations.

#### Knowledge Guided miRNA Prediction

For prediction of upregulated miRNAs a knowledge guided bioinformatics model was developed as previously described [33]. Candidate miRNAs predicted to be upregulated were identified by testing whether their TargetScanHuman 8.1 predicted targets were enriched among downregulated mRNAs from previous DGE analysis. Predicted miRNA–mRNA interactions were obtained from a raw TargetScan 8.1 context++ table (data/targetscan_context_scores.csv), restricted to human (tax_id 9606, where available), filtered by weighted context++ score (≤ −0.2), and limited to targets present in the DESeq2 universe; duplicate miRNA–gene entries were collapsed by retaining the strongest (most negative) score. For each miRNA, enrichment of its targets among downregulated genes was assessed using a right-tailed hypergeometric test and Benjamini–Hochberg correction; miRNAs with <20 targets in the universe were excluded. To prioritise miRNAs with specific downregulated targets, we computed a soft specificity score *z*_*soft*, defined as the mean of 1/degree(*g*)across downregulated targets *g*, where degree(*g*)is the number of miRNAs targeting *g*in the network. miRNAs were called candidates if they met FDR ≤ 0.01, fold enrichment ≥ 1.5, k ≥ 5 downregulated targets, and *z*_*soft* ≥ 0.2.

### Immunostaining Analysis

Cells were washed with PBS before being fixed with 3.7% PFA for 20mins at room temperature, then permeabilized with 0.3% Triton-X-100 for 10mins at room temperature and blocked with 5% donkey serum for 1 hour. Cells were then incubation with primary antibodies in PBS-T with 5% donkey serum overnight at 4 °C. Secondary antibodies were applied in PBS-T for 1.5 h at RT, counterstained with DAPI (Molecular Probes) and mounted in DAKO fluorescent mountant (Life Technologies). Samples were imaged on a Leica DMI6000b fluorescent microscope. Primary antibodies were as follows: Pax6 (1:1000, AB_528427, DSHB), MAP2 (1:500, ab92434, AbCam), REST (1:100, 07-579, Merck). Secondary antibodies were as follows: Alexa Fluor 488-conjugated donkey anti-mouse (1∶1000, A21202, Invitrogen), Alexa Fluor 647-conjugated donkey anti-chicken (1∶1000, A78952, Invitrogen). Mean fluorescence intensity measurements were taken from at least 4 biological samples.

### Sholl Analysis

Neuronal morphology was quantified using Sholl analysis to assess neurite complexity. MAP2-immunopositive neurons were imaged and individual cells were selected for analysis based on clear soma definition and isolated morphology. Concentric circles were generated at 10 µm radial increments from the centroid of the cell soma, and the number of neurite intersections at each radius was calculated. Global and fine branching patterns were analysed using the Sholl Analysis plugin in FIJI (ImageJ). For each condition, 10 neurons were analysed per differentiation across a total of six independent differentiations (n = 6).

### MultimiR-sponge design and construction

The multimiR sponges were constructed from a modified method previously reported [61]. Three empty sponge cassettes were generated by introducing a multiple cloning site containing the AgeI, NheI, SbfI and ApaI, restriction sequences, with a nonpalindromic KlfI site positioned between either the 1^st^ and 2^nd^, 2^nd^ and third, or 3^rd^ and 4^th^ restriction site. The separate MCS’ were ligated into the 3’ UTR of pcDNA3-EGFP, a gift from Doug Golenbock (Addgene plasmid # 13031), digested with XbaI and ApaI. Individual miRNA sponges were designed and tested in-silico using the miRNAsong algorithm to assess on-target and off-target binding efficiency [62]. A scramble sequence (TCATACTATATGACATCATA), denoted as Empty Vector (EV), was also designed, which targeted no known region of the human transcriptome. Individual sponges were generated as previously described [63], into one of the three MCS cassettes (A, B, C). MultimiR-sponges were then generated by digesting each of the individual sponges and ligating into a single plasmid sponge. Luciferase reporters were again designed and constructed as previously described [63]. Sponges and reporters were transfected using Lipofectamine 3000 (ThermoFisher Scientific) as per manufacturer’s instructions. For generation of retroviral multimiR-sponges, the sponge sequence was subcloned into the pBABE-puro retroviral vector, a gift from Hartmut Land & Jay Morgenstern & Bob Weinberg (Addgene plasmid # 1764). The production of amphotropic viruses and infection of target cells were described previously [64].

For gain of function studies, synthetic double stranded miRNA mimics were used (ThermoFisher Scientific). Mimics were transfected into NPCs at 50nM on day 10 of differentiation with Lipofectamine 3000 (ThermoFisher Scientific). Cells were harvested 4 days later, and RNA was isolated for analysis of miRNA expression by qRT-PCR. The mirVana negative control (ThermoFisher Scientific), was used to assess the unspecific effects of mimic treatment.

### Statistical analysis

Statistics and replicates (applied to all qPCR/Western/imaging panels unless otherwise stated): n denotes independent differentiations (biological replicates). For qRT–PCR, each biological replicate was measured in technical triplicate, and technical replicates were averaged to yield one value per biological replicate prior to statistical testing. For Western blots, each lane represents one biological replicate; band intensities were normalised to GAPDH and analysed on the biological-replicate values. Data are shown as mean ± SEM. All tests were two-sided. Significance: *P < 0.05, **P < 0.01, **P < 0.001.

## Acknowledgements

This work was supported by a PhD studentship from the GW4 Biomed MRC Doctoral Training Partnership to JSW, and from The Waterloo Foundation’s Developing Minds programme to JSW and AJH.

## Contributions

JSW, MA and AJH designed experiments, JH and AJH supervised the research project, JSW performed all wet bench experiments, wrote the code, and analysed data, and JSW and AJH wrote and edited the manuscript.

## Ethics declarations

None required

## References

1. Zhang, Y., et al., Rapid single-step induction of functional neurons from human pluripotent stem cells. Neuron, 2013. 78(5): p. 785–98.

2. Sydnor, V.J., et al., Neurodevelopment of the association cortices: Patterns, mechanisms, and implications for psychopathology. Neuron, 2021. 109(18): p. 2820–2846.

3. Ciceri, G., et al., An epigenetic barrier sets the timing of human neuronal maturation. Nature, 2024. 626(8000): p. 881–890.

4. Grove, J., et al., Identification of common genetic risk variants for autism spectrum disorder. Nature Genetics, 2019. 51(3): p. 431–444.

5. Pardiñas, A.F., et al., Common schizophrenia alleles are enriched in mutation-intolerant genes and in regions under strong background selection. Nature Genetics, 2018. 50(3): p. 381–389.

6. Chen, C.-Y., et al., The impact of rare protein coding genetic variation on adult cognitive function. Nature Genetics, 2023. 55(6): p. 927–938.

7. McCarthy, S.E., et al., De novo mutations in schizophrenia implicate chromatin remodeling and support a genetic overlap with autism and intellectual disability. Molecular Psychiatry, 2014. 19(6): p. 652–658.

8. Poisson, A., et al., Chromatin remodeling dysfunction extends the etiological spectrum of schizophrenia: a case report. BMC Medical Genetics, 2020. 21(1): p. 10.

9. Valencia, A.M., et al., Landscape of mSWI/SNF chromatin remodeling complex perturbations in neurodevelopmental disorders. Nature Genetics, 2023.

10. Mossink, B., et al., The emerging role of chromatin remodelers in neurodevelopmental disorders: a developmental perspective. Cellular and Molecular Life Sciences, 2021. 78(6): p. 2517–2563.

11. Nava, A.A. and V.A. Arboleda, The omics era: a nexus of untapped potential for Mendelian chromatinopathies. Human Genetics, 2023.

12. Harris, J.R., et al., Five years of experience in the Epigenetics and Chromatin Clinic: what have we learned and where do we go from here? Human Genetics, 2023.

13. Kaur, A., et al., Pattern Recognition of Common Multiple Congenital Malformation Syndromes with Underlying Chromatinopathy. J Pediatr Genet, 2022(EFirst).

14. Kleefstra, T., et al., Loss-of-function mutations in euchromatin histone methyl transferase 1 (EHMT1) cause the 9q34 subtelomeric deletion syndrome. The American Journal of Human Genetics, 2006. 79(2): p. 370–377.

15. Kleefstra, T., et al., Disruption of an EHMT1-associated chromatin-modification module causes intellectual disability. The American Journal of Human Genetics, 2012. 91(1): p. 73–82.

16. Bonati, M.T., et al., 9q34.3 microduplications lead to neurodevelopmental disorders through EHMT1 overexpression. neurogenetics, 2019. 20(3): p. 145–154.

17. Benevento, M., et al., Histone Methylation by the Kleefstra Syndrome Protein EHMT1 Mediates Homeostatic Synaptic Scaling. Neuron, 2016. 91(2): p. 341–355.

18. Frega, M., et al., Neuronal network dysfunction in a model for Kleefstra syndrome mediated by enhanced NMDAR signaling. Nature communications, 2019. 10(1): p. 4928–4928.

19. Balemans, M.C., et al., Reduced exploration, increased anxiety, and altered social behavior: Autistic-like features of euchromatin histone methyltransferase 1 heterozygous knockout mice. Behav Brain Res, 2010. 208(1): p. 47–55.

20. Alsaqati, M., et al., NRSF/REST lies at the intersection between epigenetic regulation, miRNA-mediated gene control and neurodevelopmental pathways associated with Intellectual disability (ID) and Schizophrenia. Translational Psychiatry, 2022. 12(1): p. 438.

21. Wu, Y.E., et al., Genome-wide, integrative analysis implicates microRNA dysregulation in autism spectrum disorder. Nature Neuroscience, 2016. 19(11): p. 1463–1476.

22. Sotoudeh Anvari, M., et al., Identification of microRNAs associated with human fragile X syndrome using next-generation sequencing. Scientific Reports, 2022. 12(1): p. 5011.

23. Rey, R., et al., Widespread transcriptional disruption of the microRNA biogenesis machinery in brain and peripheral tissues of individuals with schizophrenia. Translational Psychiatry, 2020. 10(1): p. 376.

24. Fear, V.S., et al., CRISPR single base editing, neuronal disease modelling and functional genomics for genetic variant analysis: pipeline validation using Kleefstra syndrome EHMT1 haploinsufficiency. Stem Cell Research & Therapy, 2022. 13(1): p. 69.

25. Gennady, K., et al., Fast gene set enrichment analysis. bioRxiv, 2021: p. 060012.

26. Lake, B.B., et al., Integrative single-cell analysis of transcriptional and epigenetic states in the human adult brain. Nature Biotechnology, 2018. 36(1): p. 70–80.

27. Piñero, J., et al., DisGeNET: a comprehensive platform integrating information on human disease-associated genes and variants. Nucleic Acids Research, 2017. 45(D1): p. D833–D839.

28. Quartier, A., et al., Novel mutations in NLGN3 causing autism spectrum disorder and cognitive impairment. Human Mutation, 2019. 40(11): p. 2021–2032.

29. Tomaselli, P.J., et al., A de novo dominant mutation in KIF1A associated with axonal neuropathy, spasticity and autism spectrum disorder. Journal of the Peripheral Nervous System, 2017. 22(4): p. 460–463.

30. Kaneko, M., T. Rosser, and G. Raca, Dilated cardiomyopathy in a patient with autosomal dominant EEF1A2-related neurodevelopmental disorder. European Journal of Medical Genetics, 2021. 64(1): p. 104121.

31. Tripathy, R., et al., Mutations in MAST1 Cause Mega-Corpus-Callosum Syndrome with Cerebellar Hypoplasia and Cortical Malformations. Neuron, 2018. 100(6): p. 1354–1368.e5.

32. Cooper, G.M., et al., A copy number variation morbidity map of developmental delay. Nat Genet, 2011. 43(9): p. 838–46.

33. Shen, L., et al., Knowledge-Guided Bioinformatics Model for Identifying Autism Spectrum Disorder Diagnostic MicroRNA Biomarkers. Scientific Reports, 2016. 6(1): p. 39663.

34. Zhang, W., et al., Identification of candidate miRNA biomarkers from miRNA regulatory network with application to prostate cancer. J Transl Med, 2014. 12: p. 66.

35. Wu, J. and X. Xie, Comparative sequence analysis reveals an intricate network among REST, CREBand miRNA in mediating neuronal gene expression. Genome biology, 2006. 7(9): p. 1–14.

36. Richner, M., et al., MicroRNA-based conversion of human fibroblasts into striatal medium spiny neurons. Nature Protocols, 2015. 10(10): p. 1543–1555.

37. Wu, S., et al., Multiple microRNAs modulate p21Cip1/Waf1 expression by directly targeting its 3ʹ untranslated region. Oncogene, 2010. 29(15): p. 2302–2308.

38. Ebert, M.S., J.R. Neilson, and P.A. Sharp, MicroRNA sponges: competitive inhibitors of small RNAs in mammalian cells. Nature Methods, 2007. 4(9): p. 721–726.

39. Lee, S.W., et al., MicroRNAs Overcome Cell Fate Barrier by Reducing EZH2-Controlled REST Stability during Neuronal Conversion of Human Adult Fibroblasts. Dev Cell, 2018. 46(1): p. 73–84.e7.

40. Chen, R., et al., Homeodomain protein Six4 prevents the generation of supernumerary Drosophila type II neuroblasts and premature differentiation of intermediate neural progenitors. PLoS Genet, 2021. 17(2): p. e1009371.

41. Blackwood, C.A., Jagged1 is Essential for Radial Glial Maintenance in the Cortical Proliferative Zone. Neuroscience, 2019. 413: p. 230–238.

42. Latremoliere, A., et al., Neuronal-Specific TUBB3 Is Not Required for Normal Neuronal Function but Is Essential for Timely Axon Regeneration. Cell reports, 2018. 24(7): p. 1865–1879.e9.

43. Peláez, N. and R.W. Carthew, Biological robustness and the role of microRNAs: a network perspective. Curr Top Dev Biol, 2012. 99: p. 237–55.

44. Stappert, L., et al., MicroRNA-based promotion of human neuronal differentiation and subtype specification. PloS one, 2013. 8(3): p. e59011.

45. Conaco, C., et al., Reciprocal actions of REST and a microRNA promote neuronal identity. Proceedings of the National Academy of Sciences, 2006. 103(7): p. 2422–2427.

46. Cao, X., S.L. Pfaff, and F.H. Gage, A functional study of miR-124 in the developing neural tube. Genes Dev, 2007. 21(5): p. 531–6.

47. Kowalczyk, A., et al., The critical role of cyclin D2 in adult neurogenesis. J Cell Biol, 2004. 167(2): p. 209–13.

48. Gebhardt, M.L., et al., Similarity in targets with REST points to neural and glioblastoma related miRNAs. Nucleic Acids Research, 2014. 42(9): p. 5436–5446.

49. Sauer, M., et al., The miR-26 family regulates neural differentiation-associated microRNAs and mRNAs by directly targeting REST. J Cell Sci, 2021. 134(12).

50. Diener, C., et al., Outside the limit: questioning the distance restrictions for cooperative miRNA binding sites. Cell Mol Biol Lett, 2023. 28(1): p. 8.

51. Flamand, M.N., et al., A non-canonical site reveals the cooperative mechanisms of microRNA-mediated silencing. Nucleic Acids Research, 2017. 45(12): p. 7212–7225.

52. Petanjek, Z., et al., The Protracted Maturation of Associative Layer IIIC Pyramidal Neurons in the Human Prefrontal Cortex During Childhood: A Major Role in Cognitive Development and Selective Alteration in Autism. Front Psychiatry, 2019. 10: p. 122.

53. Sehovic, E., et al., Identification of developmental disorders including autism spectrum disorder using salivary miRNAs in children from Bosnia and Herzegovina. PLoS One, 2020. 15(4): p. e0232351.

54. Mor, M., et al., Hypomethylation of miR-142 promoter and upregulation of microRNAs that target the oxytocin receptor gene in the autism prefrontal cortex. Molecular autism, 2015. 6(1): p. 1–11.

55. Perkins, D.O., et al., microRNA expression in the prefrontal cortex of individuals with schizophrenia and schizoaffective disorder. Genome biology, 2007. 8(2): p. 1–11.

56. Perkins, D.O., C. Jeffries, and P. Sullivan, Expanding the ‘central dogma’: the regulatory role of nonprotein coding genes and implications for the genetic liability to schizophrenia. Mol Psychiatry, 2005. 10(1): p. 69–78.

57. Li, Y., et al., Aberrant expression of miR-153 is associated with overexpression of hypoxia-inducible factor-1α in refractory epilepsy. Scientific Reports, 2016. 6(1): p. 32091.

58. You, Y.-H., et al., MicroRNA-153 promotes brain-derived neurotrophic factor and hippocampal neuron proliferation to alleviate autism symptoms through inhibition of JAK-STAT pathway by LEPR. Bioscience reports, 2019. 39(6): p. BSR20181904.

59. Chambers, S.M., et al., Highly efficient neural conversion of human ES and iPS cells by dual inhibition of SMAD signaling. Nature Biotechnology, 2009. 27(3): p. 275–280.

60. Piñero, J., et al., The DisGeNET cytoscape app: Exploring and visualizing disease genomics data. Computational and Structural Biotechnology Journal, 2021. 19: p. 2960–2967.

61. Kluiver, J., et al., Generation of miRNA sponge constructs. Methods, 2012. 58(2): p. 113–7.

62. Barta, T., L. Peskova, and A. Hampl, miRNAsong: a web-based tool for generation and testing of miRNA sponge constructs in silico. Scientific reports, 2016. 6(1): p. 1–8.

63. Kluiver, J., et al., Generation of miRNA sponge constructs. Methods, 2012. 58(2): p. 113–117.

64. Stewart, S.A., et al., Lentivirus-delivered stable gene silencing by RNAi in primary cells. Rna, 2003. 9(4): p. 493–501.

