## Supplementary material for "Control of timing and directionality during neurodifferentiation of human induced Pluripotent Stem Cells (iPSC) via miRNA-mediated feedback and feedforward loops": Fig S1

**Supplemental Figures**

**Figure S1: Permutation testing of observed RE1 taregt/upregulated gene overlap**

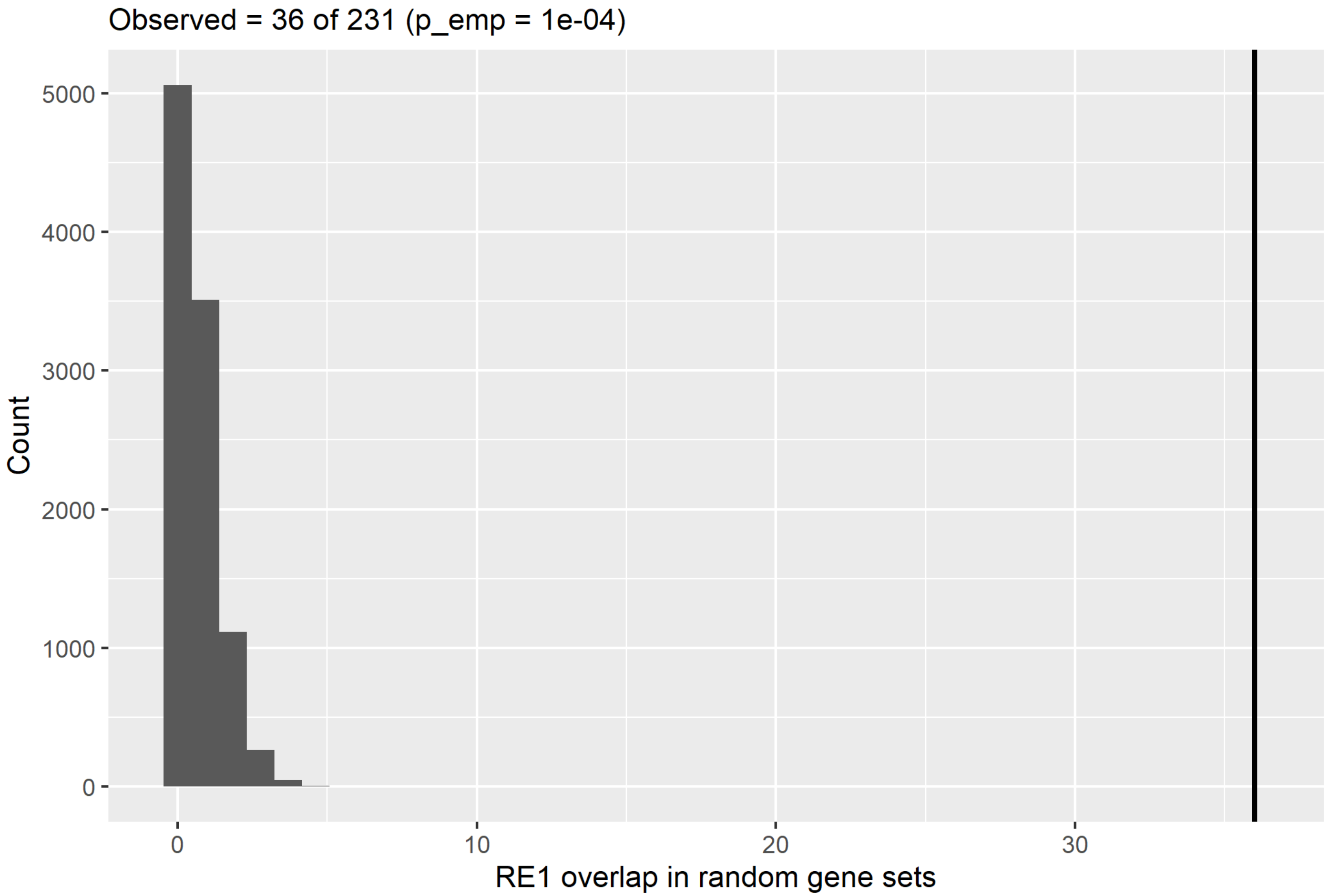


RE1 enrichment was assessed by permutation testing using 10,000 random gene sets sampled from expressed genes (baseMean > 10); empirical P values were computed as (r+1)/(n+1).
