## Supplementary material for "Control of timing and directionality during neurodifferentiation of human induced Pluripotent Stem Cells (iPSC) via miRNA-mediated feedback and feedforward loops": Fig S2

**Figure S2: Enrichment analysis of EHMT1+/- neuronal cells**

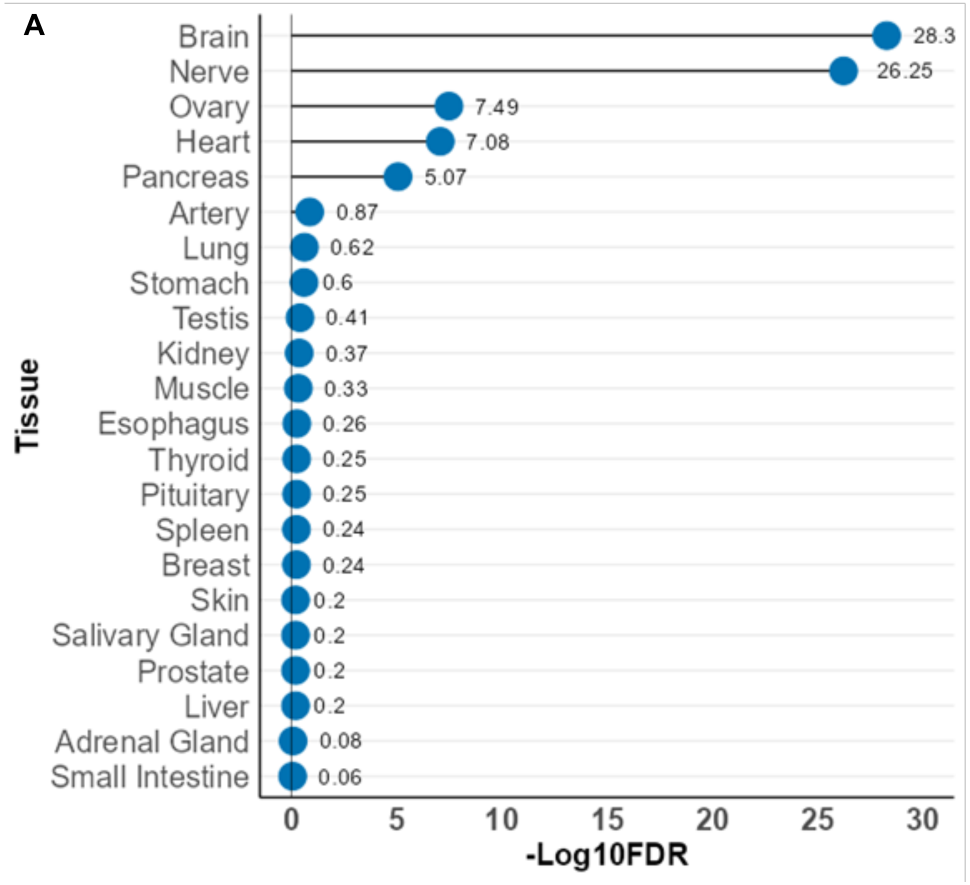


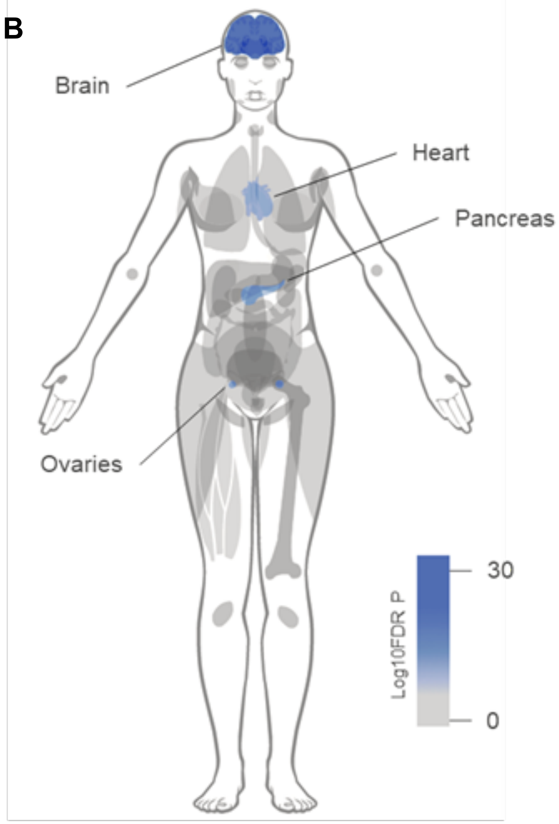


**A:** Tissue enrichment analysis using gene expression per tissue based on GTEx RNA-seq data for general tissues

**B:** Schematic body map of significantly enriched tissues.
