## Supplementary material for "Control of timing and directionality during neurodifferentiation of human induced Pluripotent Stem Cells (iPSC) via miRNA-mediated feedback and feedforward loops": Fig S3

**Figure S3: Analysis of REST protein levels**

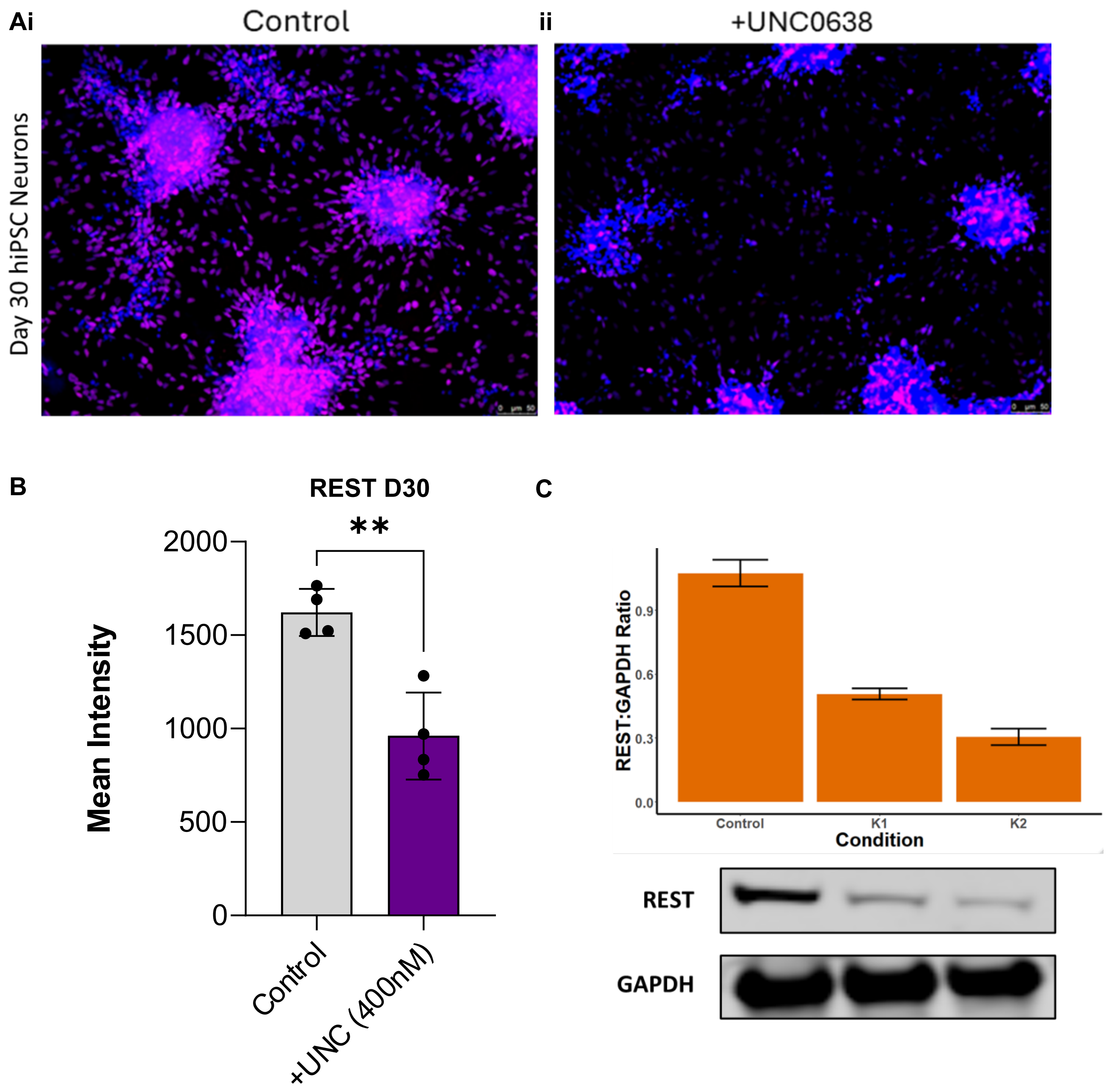


**A:** Analysis of REST (Purple) protein expression in control and UNC0638 treated (400nM) cells. Images were acquired using Leica DMI6000b fluorescent microscope. Scale bar, 50µM. **B:** Quantification of protein expression as REST mean fluorescence intensity, with DAPI nuclear counterstain. (** $p<0.01$). Figure B shows summary data from 4 samples across 4 differentiations. **C:** Analysis of REST protein expression in hiPSCs derived from two independent KS patients (KS1, KS2). REST expression was decreased in both patient lines as compared to control line, n=6
