## Supplementary material for "Control of timing and directionality during neurodifferentiation of human induced Pluripotent Stem Cells (iPSC) via miRNA-mediated feedback and feedforward loops": Fig S4

**Figure S4: ChIP-qPCR analysis of H3K9me2 marks on miRNA targets**

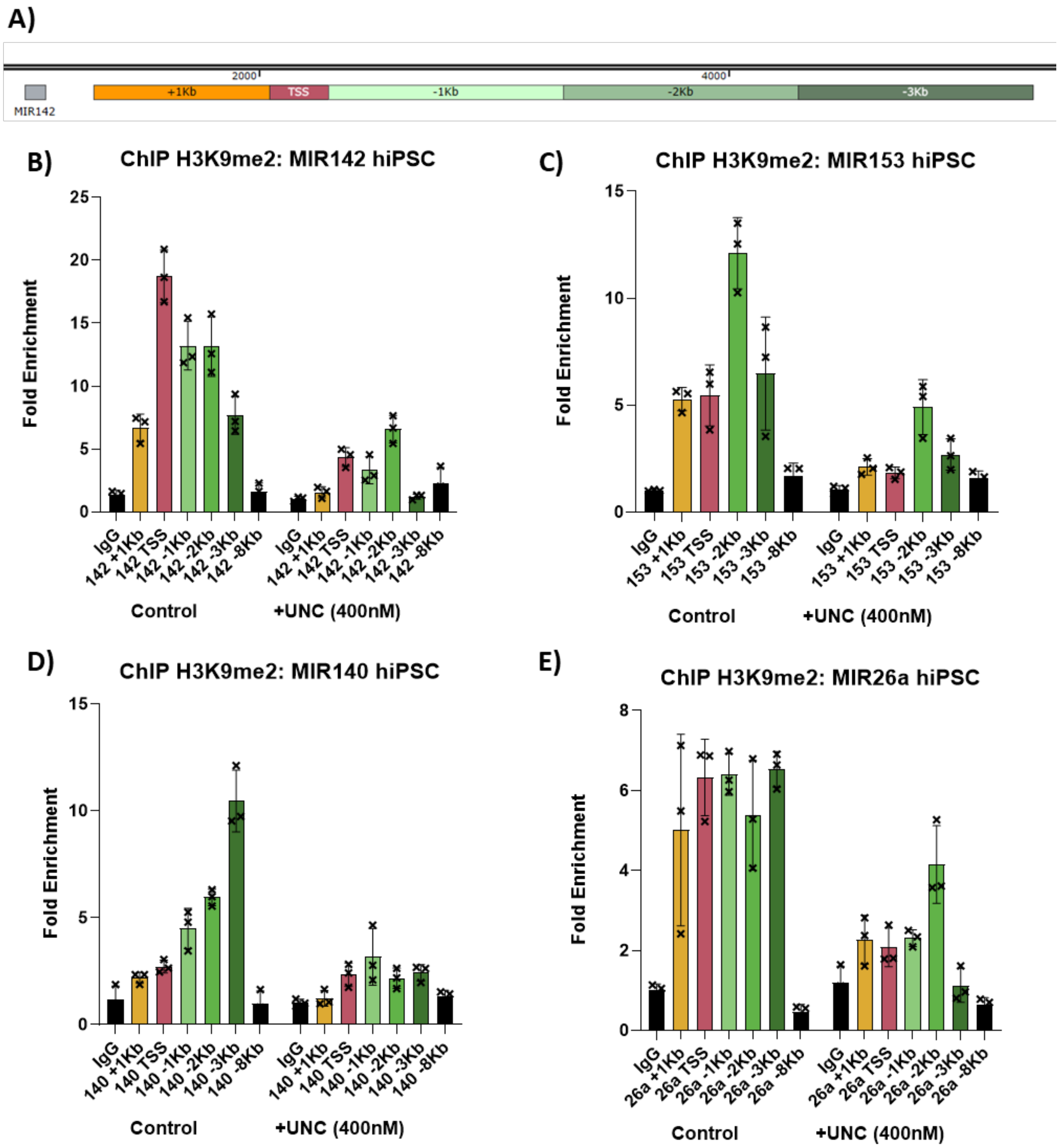


**A:** Schematic of regions in the MIR142 gene analysed by qPCR. Primer pairs were designed at the transcription start site (TSS), 1Kb downstream of the TSS, or up to -3Kb upstream of the TSS at 1Kb intervals. B-E Analysis of miRNAs in hiPSCs treated with 400nM UNC0638 as compared to vehicle control. Mean fold change over vehicle control, n = 3 independent experiments **B:** miR142 **C:** miR-153, successful primers were not possible for -1Kb downstream of the TSS, **D:** miR140, **E:** miR26a. Data were presented as Mean±SEM, from n = 3 independent experiments.
