## Supplementary material for "Control of timing and directionality during neurodifferentiation of human induced Pluripotent Stem Cells (iPSC) via miRNA-mediated feedback and feedforward loops": Fig S5

**Figure S5: MultimiR-sponges can effectively suppress miRNAs**

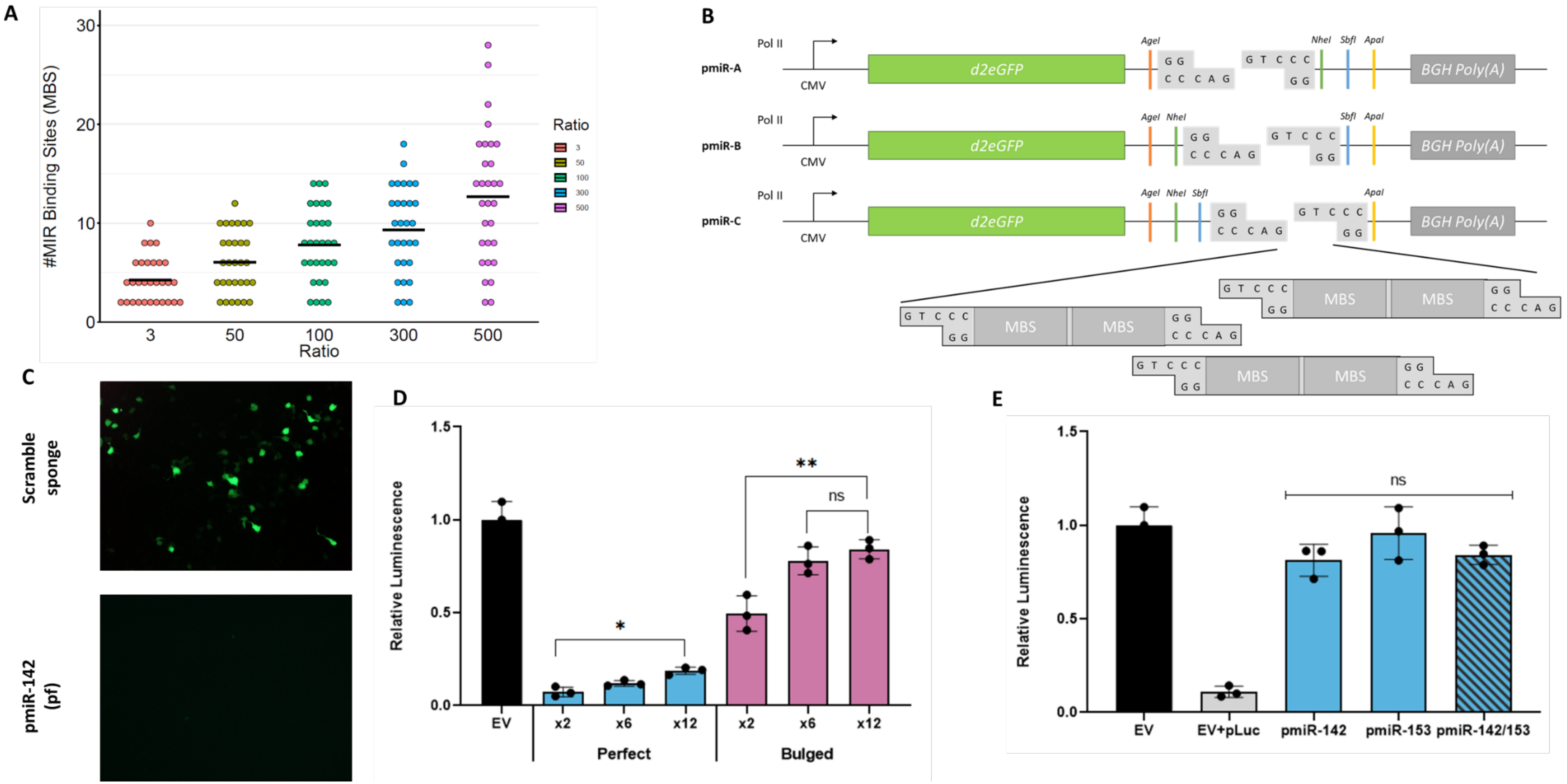


**A:** Analysis of microRNA binding site inserts proportional to insert ratio. By increasing the insert cloning ratio, the number of binding sites can be significantly increased. **B:** Overview of the multimiR sponge cloning approach. Three vector libraries allow for subcloning of individual miR sponges to generate the combined multimiR-sponges. **C, D:** Efficacy analysis of individual sponge inserts using luciferase reporters. **(C)** Immunofluorescence imaging of perfect sponges demonstrates degradation of the miR-sponge as indicated by a lack of eGFP expression. **(D)** Luciferase reporter analysis supported the finding that perfect sponges were degraded, whilst bulged sequence remained intact and were capable of miR repression. **E:** Luciferase reporter analysis demonstrated that individual sponges were able to repress specific miRNAs, whilst the multimiR-sponge showed no significant drop in efficacy when targeting miR's simultaneously.
