## Supplementary material for "Control of timing and directionality during neurodifferentiation of human induced Pluripotent Stem Cells (iPSC) via miRNA-mediated feedback and feedforward loops": Fig S6

**Figure S6: Temporal effects of EHMT1 loss**

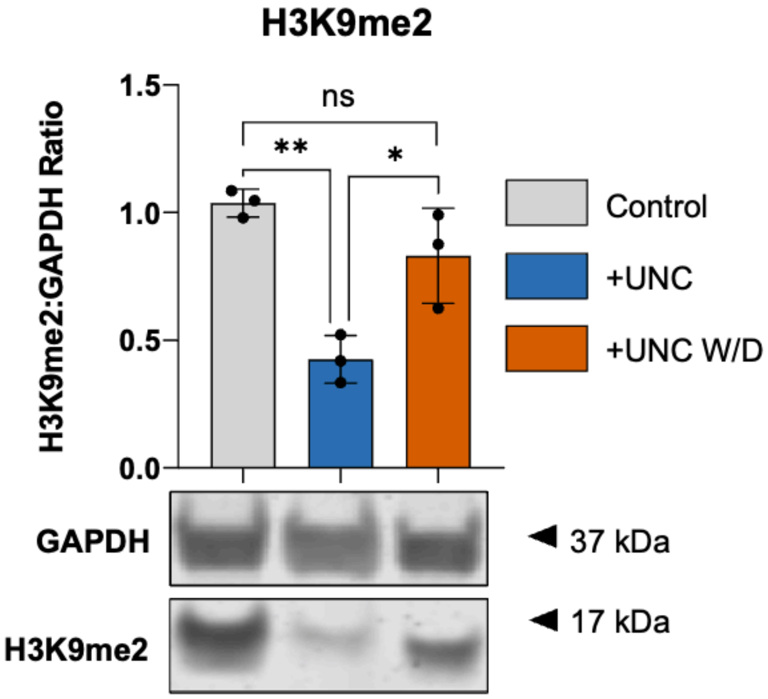


Western blot analysis of H3K9me2 protein expression in hiPSC derived neurons at day 30 following treatment with UNC0638 (UNC) from days 0-10, as compared to vehicle control. Withdrawal (W/D) of UNC0638 restored H3K9me2 protein to control levels.
