## Supplementary material for "Control of timing and directionality during neurodifferentiation of human induced Pluripotent Stem Cells (iPSC) via miRNA-mediated feedback and feedforward loops": Fig S7

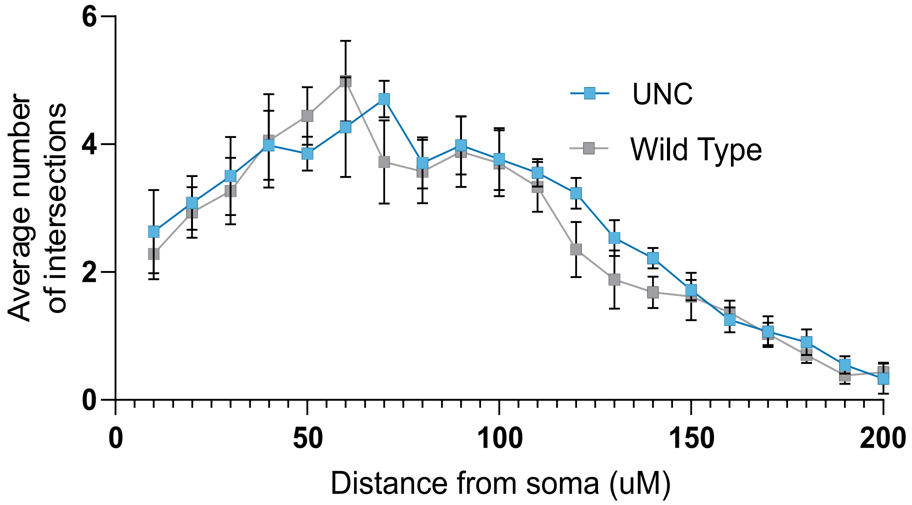


**Figure S7: Quantification of neurite branching complexity in induced neurons.**

Sholl analysis, showing the average number of intersections as a function of distance from the soma (µm). no difference in neurite complexity is seen between UNC treated to wild-type controls. Data were presented as Mean ± SEM from n = 3 independent experiments.
