## Supplementary material for "Control of timing and directionality during neurodifferentiation of human induced Pluripotent Stem Cells (iPSC) via miRNA-mediated feedback and feedforward loops": Table S1

Primers for mature miRNA quantitation.

| Target | Sequence (5’-3’) |
|---|---|
| **Universal** | GAATCGAGCACCAGTTACGC |
| **miR-142-3p** | TGTAGTGTTTCCTACTTTATGGA |
| **miR-142-5p** | GCAGCATAAAGTAGAAAGCACT |
| **miR-153-3p** | TTGCATAGTCACAAAAGTGATC |
| **miR-153-5p** | GCAGTTGCATAGTCACAAAAGT |
| **miR-26a-3p** | TTATAATACAACCTGATAAGTG |
| **miR-26a-5p** | TTCAAGTAATCCAGGATAGGCT |
| **miR-26b-3p** | CCTGTTCTCCATTACTTGGCT |
| **miR-26b-5p** | GGCTTCAAGTAATTCAGGATAGG |
| **miR-27b-3p** | TTCACAGTGGCTAAGTTCTGC |
| **miR-27b-5p** | AGAGCTTAGCTGATTGGTGAAC |
| **miR-320a-3p** | AAAAGCTGGGTTGAGAGGGCGA |
| **miR-320a-5p** | GCCTTCTCTTCCCGGTTCTTCC |
| **miR-340-3p** | TCCGTCTCAGTTACTTTATAGC |
| **miR-340-5p** | TTATAAAGCAATGAGACTGATT |
| **miR-548f-3p** | AAAAACTGTAATTACTTTT |
| **miR-548f-5p** | GCAGTGCAAAAGTAATCACAGT |
| **miR-653-3p** | TTCACTGGAGTTTGTTTCAATA |
| **miR-653-5p** | GTGTTGAAACAATCTCTACTG |

| Target | Sequence (5’-3’) |
|---|---|
| **miR-101-3p** | GCTACAGTACTGTGATAACTG |
| **miR-101-5p** | CAGTTATCACAGTGCTGATGCT |
| **miR-10b-3p** | ACAGATTCGATTCTAGGGGAAT |
| **miR-124-3p** | TAAGGCACGCGGTGAATGCC |
| **miR-124-5p** | CGTGTTCACAGCGGACCTTGAT |
| **miR-135b-3p** | ATGTAGGGCTAAAAGCCATGGG |
| **miR-135b-5p** | TATGGCTTTTCATTCCTATGTGA |
| **miR-140-3p** | TACCACAGGGTAGAACCACGG |
| **miR-140-5p** | CAGTGGTTTTACCCTATGGTAG |
| **miR-144-3p** | GCTACAGTATAGATGATGTACT |
| **miR-144-5p** | GCGGATATCATCATATACTGTA |
| **miR-218-3p** | ATGGTTCCGTCAAGCACCATGG |
| **miR-218-5p** | TTGTGCTTGATCTAACCATGT |
| **miR-27a-3p** | TTCACAGTGGCTAAGTTCCGC |
| **miR-340-5p** | GCAGTTATAAAGCAATGAGACTGA |
| **miR-6835-3p** | AAAAGCACTTTTCTGTCTCCCAG |
| **miR-6835-5p** | AGGGGGTAGAAAGTGGCTGAAG |
| **miR-9-3p** | ATAAAGCTAGATAACCGAAAGT |
| **miR-9-5p** | TCTTTGGTTATCTAGCTGTATGA |

| Gene | Forward Sequence (5’-3’) | Reverse Sequence (5’-3’) |
|---|---|---|
| **ACTA1** | GGCATTCACGAGACCACCTAC | CGACATGACGTTGTTGGCATAC |
| **ACTL6B** | GCGCTGGTCTTTGACATTGG | CCATTCTTGAGGGGCGACAT |
| **AP3B1** | GAAGCGGATTGTTGGGATGAT | TCAGCATATCGAACCAGGTAAAC |
| **ASCL1** | CGCGGCCAACAAGAAGATG | CGACGAGTAGGATGAGACCG |
| **CALB1** | TGGCATCGGAAGAGCAGCAG | TGACGGAAGTGGTTACCTGGAAG |
| **CDK6** | CCAGATGGCTCTAACCTCAGT | AACTTCCACGAAAAAGAGGCTT |
| **CITED2** | CCTAATGGGCGAGCACATACA | GGGGTAGGGGTGATGGTTGA |
| **CTDSPL** | CCACCAGCTAAGTACCTTCTTCC | GGCCGCTTCAGCACATACA |
| **ETS1** | GATAGTTGTGATCGCCTCACC | GTCCTCTGAGTCGAAGCTGTC |
| **FREM2** | CCTGCATGACCTGGTGTTG | GCCAGTGCGTCGTTGTCTA |
| **GAPDH** | ACCACAGTCATGCCATCAC | TCCACCACCCTGTTGCTGTA |
| **JAG1** | GTCCATGCAGAACGTGAACG | GCGGGACTGATACTCCTTGA |
| **ITGB1** | CCGCGCGGAAAAGATGAAT | ATGTCATCTGGAGGGCAACC |
| **KIF20A** | TGCTGTCCGATGACGATGTC | AGGTTCTTGCGTACCACAGAC |
| **LAMC1** | GGACTCCGCCCGAGGAATA | ACTTGAGACGCACATAGGTGA |
| **MAP2** | CTGCTTTACAGGGTAGCACAA | TTGAGTATGGCAAACGGTCTG |
| **NCAM1** | GGCATTTACAAGTGTGTGGTTAC | TTGGCGCATTCTTGAACATGA |
| **NRXN3** | AGTGGTGGGCTTATCCTCTAC | CCCTGTTCTATGTGAAGCTGGA |
| **ONECUT2** | GGAATCCAAAACCGTGGAGTAA | CTCTTTGCGTTTGCACGCTG |
| **PAX6** | GTGTCCAACGGATGTGTGAG | CTAGCCAGGTTGCGAAGAAC |
| **PHF6** | AGAGGCACGAAGCTGATGTG | AGTGGTAGTGGTATGTCCTGTG |
| **PRRX1** | TGATGCTTTTGTGCGAGAAGA | AGGGAAGCGTTTTTATTGGCT |
| **REST** | GCCGCACCTCAGCTTATTATG | CCGGCATCAGTTCTGCCAT |
| **ROCK1** | AACATGCTGCTGGATAAATCTGG | TGTATCACATCGTACCATGCCT |
| **SIX4** | AGCAGCTCTGGTACAAGGC | CTTGAAACAATACACCGTCTCCT |
| **SNORD48** | AGTGATGATGACCCCAGGTAACTC | GGTCAGAGCGCTGCGGTG |
| **UNC5C** | TGGGACTGGGATACTTGCTG | ACAGTACAGGTTCACAGGCTTAT |
