## Supplementary material for "Control of timing and directionality during neurodifferentiation of human induced Pluripotent Stem Cells (iPSC) via miRNA-mediated feedback and feedforward loops": Table S2

REST Context++ and PCt scores from TargetScan of miRNAs upregulated in hiPSCs treated with UNC0638.

| miRNA | Log2FC | Context++ Score | PCt |
|---|---|---|---|
| **miR-142-3p** | 13.38 | -0.09 | 0.36 |
| **miR-153-3p** | 8.55 | -0.28 | 0.82 |
| **miR-140-5p** | 6.51 | -0.38 | 0.33 |
| **mIR-26a-5p** | 5.50 | -0.20 | 0.57 |
